# A Phytosociological Classification of the Peruvian Vegetation

**DOI:** 10.1101/2021.03.17.435755

**Authors:** Antonio Galán-de-Mera, José Campos-de-la-Cruz, Eliana Linares-Perea, Juan Montoya-Quino, Iván Torres-Marquina, José Alfredo Vicente-Orellana

## Abstract

The present work is an update of the syntaxonomic scheme of Peru, with the results obtained to present. We currently recognize 218 phytosociological associations, of which 11 are novelties: *Ageratino azangaroensis-Polylepidetum incarum* ass. nov., *Alchemillo pinnatae-Stipetum hans-meyeri* ass. nov., *Barnadesio dombeyanae-Polylepidetum racemosae* ass. nov., *Eclipto prostratae-Echinochloetum crus-pavonis* ass. nov., *Geoffroeetum decorticantis* ass. nov., *Junellio arequipensis-Proustietum cuneifoliae* ass. nov., *Lemnetum aequinoctialis* ass. nov., *Mutisio cochabambensis-Polylepidetum incarum* ass. nov., *Poo glaberrimae-Calamagrostietum eminentis* ass. nov., *Sarcocornietum andinae* ass. nov., and *Schino mollis-Prosopidetum calderensis* ass. nov. These 218 associations belong to 34 phytosociological classes, four of which are described here for the first time: *Limoselletea australis* cl. nov., *Mimulo glabrati-Plantaginetea australis* cl. nov., *Tillandsietea latifolio-landbeckii* cl. nov., *Woodsio montevidensis-Cheilanthetea pruinatae* cl. nov. In addition, 5 new orders (*Alternanthero pungentis-Cynodontetalia dactylonis* ord. nov., *Coreopsietalia fasciculatae* ord. nov., *Echinopsio schoenii-Proustetalia cuneifoliae* ord. nov., *Mimulo glabrati-Plantaginetalia australis* ord. nov., *Tillandsietalia latifolio-landbeckii* ord. nov.), and 7 new alliances (*Azollo filiculoidis-Lemnion gibbae* all. nov., *Ditricho submerse-Isoetion lechleri* all. nov., *Echinopsio schoenii-Proustion cuneifoliae* all. nov., *Haageocerion acranthi* all. nov., *Lepidio chichicarae-Cynodontion dactylonis* all. nov., *Schoenoplection tatorae* all. nov., and *Tillandsion purpureo-latifoliae* all. nov.) are also described.

## Introduction

We have been studying the vegetation of Peru using the phytosociological method of Braun-Blanquet (1932) since 1987, taking as a basis variuos works published in the area, specifically those by Cuatrecasas in Colombia (Cuatrecasas 1934), and Rivas Martínez & Tovar (1982) in Peru. We began with the study of the bioclimatic belts of Peru in the Natural History Museum of the Universidad Nacional Mayor de San Marcos, obtaining some conclusive observations regarding the parallelism between the intervals of the thermicity index and precipitation with plant communities (Galán de Mera et al. 2017). The problematic political situation in Peru during the 1980s forced us to abandon our work, which we continued in the surrounding countries, including Chile, Bolivia and Paraguay until the end of the 1990s. We later returned to Peru to continue our studies on vegetation, now with the support of the Universidad CEU San Pablo Madrid, and the Agencia Española de Cooperación Internacional y Desarrollo. We started by following the investigations of Gutte and Müller (Müller 1985a), Bolòs et al. (1991) and the pioneering paper of Navarro in the Puna of Bolivia (Navarro 1993). We prepared an initial study on the Peruvian coast and the cactus communities of the occidental slopes of the Andes, and we completed a first syntaxonomical scheme at the alliance level (Galán de Mera et al. 2002), where previous research on the Amazonia was included. After, we continued with the high mountains of southern Peru (Galán de Mera et al. 2003), describing several associations of central Peru (Galán de Mera et al. 2004), and a first assay on the syntaxomonomical scheme of Caribbean and South American vegetation (Galán de Mera 2005, Galán de Mera & Vicente Orellana 2006). This assay resulted from the comparison between the known vegetation of Peru with other regions of America, such as the French Antilles (Foucault 1978, 1981), Cuba (Borhidi 1991, 1996), Venezuela (Castroviejo & López 1985, Galán de Mera et al. 2006), Colombia (Rangel & Franco 1985), northern Chile (Luebert & Gajardo 2005), Patagonia (Boelcke et al. 1985), Bolivia (Seibert & Mehofer 1991, 1992, Seibert 1993), and other territories.

In 2008, we initiated contacts with the Universidad Nacional de San Agustín de Arequipa, which allowed us to focus on the vegetation of other regions of southern Peru, such as the departments of Arequipa and Ica, and on the eastern side, the department of Puno, where we started the phytosociological study of Andean rainforests. In parallel, we also extended in the departments of Lima, Junín and Ancash to address the vegetation linked to cryogenic processes (Galán de Mera et al. 2014). This last study culminated with the description of the new genus *Anticona* (Linares Perea et al. 2014) with one population in the Department of Lima, after the description of *Werneria glareophila* by Cuatrecasas (1970) with plants from Huancavelica.

However, 2013 was the year of the discovery of northern Peru derived from our contact with the herbarium of the Universidad Nacional de Cajamarca, whose head, Prof. Isidoro Sánchez Vega, accompanied us on some expeditions, contributing to a new research in the vegetation of the Páramos of Cajamarca and their relationship with the Ecuadorians (Galán de Mera et al. 2015), and of the western slopes of northern Peru (Galán de Mera et al. 2016). At that time, the Universidad Privada Antonio Guillermo Urrelo of Cajamarca was our logistic support through Prof. Iván Torres Marquina. After the death of Prof. Sanchez, we started studies in the Amazon and Cajamarca rainforests. All the rainforests investigated in Peru were gathered together in a recent publication (Galán de Mera et al. 2020).

The classification that we now propose, is presented using the same methodology as in other contemporary works in other parts of the world, such as Venezuelan savannas (Galán de Mera 2014), Central America (García Fuentes et al. 2014), Argentina (Martínez Carretero et al. 2016), Europe (Mucina et al. 2016), Dominican Republic (Cano Ortiz et al. 2020), American mangroves (García Fuentes et al. 2020), and the Colombian páramo (Pinto Zárate et al. in prep.).

## Material and methods

### The phytosociological method

The phytosociological method allows us to delve deeply into the analysis and classification of ecosystems and additionally, into ecological, dynamic, and geographic comparisons with other disjunct areas. These results cannot be achieved with a purely physiognomic method. The analysis of the vegetation requires the realization of a large number of phytosociological plots, which implies a deep knowledge of the flora of the territories. Then, from the variation of vegetation in disjunct areas, we can define or identify a complete hierarchy of syntaxons from the level of association to phytosociological class. Therefore, with the description of each association in a territory we are giving an image of its floristic, dynamic, chorological and historical characteristics (Braun-Blanquet 1932, Géhu & Rivas-Martínez 1981, Dierschke 1994, Rivas-Martínez 2005, Dengler et al. 2008).

A phytosociological plot is a list of species where each one is accompanied by an abundance-dominance index, which means an estimation of the percentage of coverage for each species in a determined biotope or geomorphological space (Fig. 1). The values are: +: Scarce plants, of weak coverage (< 1%), 1: Abundant plants, but weak coverage (1-5%), 2: Plants abundant but covering at least 1/20 of the surface (5-25%), 3: Plants in variable number, but covering 1/4 to 1/2 of the surface (25-50%), 4: Plants in variable number, but covering 1/2 to 3/4 of the surface (50-75%), 5: Plants in a variable number, but covering more than 3/4 of the surface (>75%).

**Figure 1.**
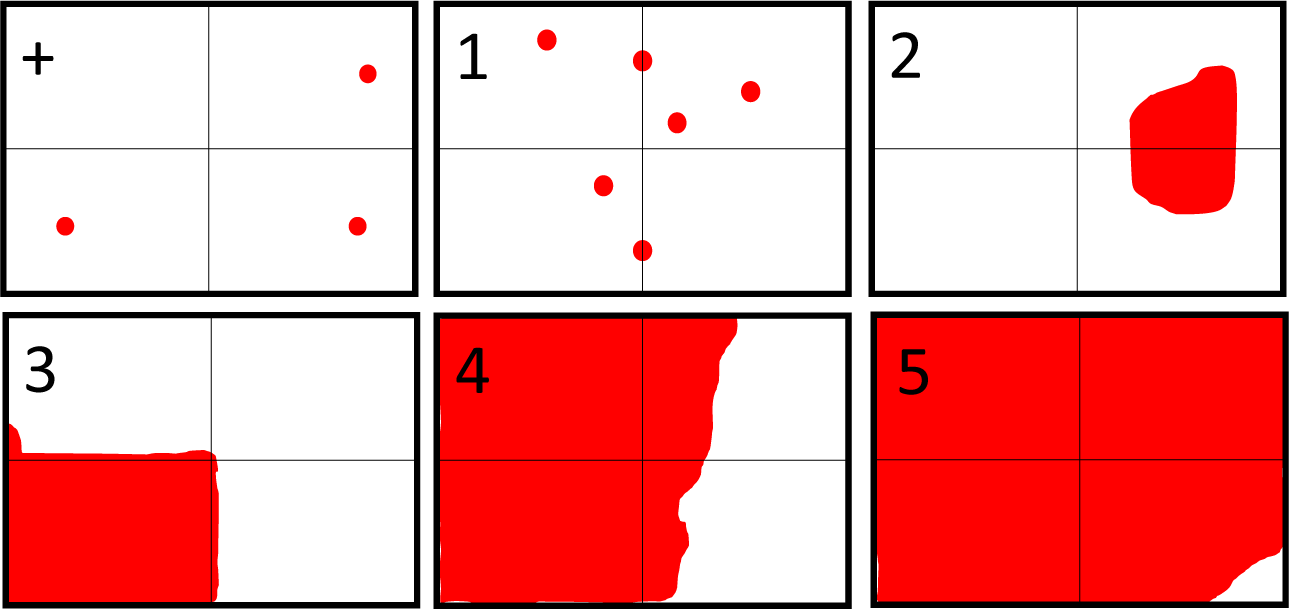
Estimation of the percentage of coverage for each species in a determined biotope or geomorphological space.

A phytosociological plot is made up of a minimum area that shares all the species of the biotope. With numerous plots carried out in a geographic area, we are able to build an association, which will be differentiated from another by a distinct floristic composition.

Among the plants of a plot, we can differentiate between characteristic, differential, and companion plants. Characteristics are those plants that due to their fidelity are exclusively linked to a given syntaxon (association, alliance, order or class). Differential plants are those that, without necessarily being characteristic, can help in the local understanding of phytosociological units. Companions are plants present in numerous groupings that are usually in contact with the one being studying. Often, companions can be used as differentials of an association by providing precise geographic or ecological information.

The associations (ending *’etum’*) are seated on a biotope or geomorphological site and have a specific geographic distribution. These are grouped into alliances (ending *’ion’*), alliances into orders (ending *’etalia’*), and orders into classes (ending *’etea’*). Associations, alliances, orders, and classes are syntaxons, and each of them, such as botanical or zoological taxa, has a specific nomenclature regulated by the International Code of Phytosociological Nomenclature (Theurillat et al. 2021).

Although the associations are located in their own geomorphological area, they also have a parallelism with bioclimatic belts.

### Bioclimatic belts in Peru

Bioclimatic belts are based on the thermicity index (It) and bioindicators, which are plants and associations (Rivas-Martínez et al. 1999). The thermicity index is a mathematical expression with different temperature values in degrees centigrade: It = (T+M+m) 10 [T: Average annual temperature, M: Average maximum temperature in the coldest month, m: Average minimum temperature of the coldest month].

At present, we are able to distinguish six bioclimatic belts throughout Peru (Galán de Mera et al. 2017; Fig. 2): Infratropical (It > 690), thermotropical (It = 490 to 690), mesotropical (It = 320 to 490), supratropical (It = 160 to 320), orotropical (It = 50 to 160), and cryorotropical (It = < 50).

**Figure 2.**
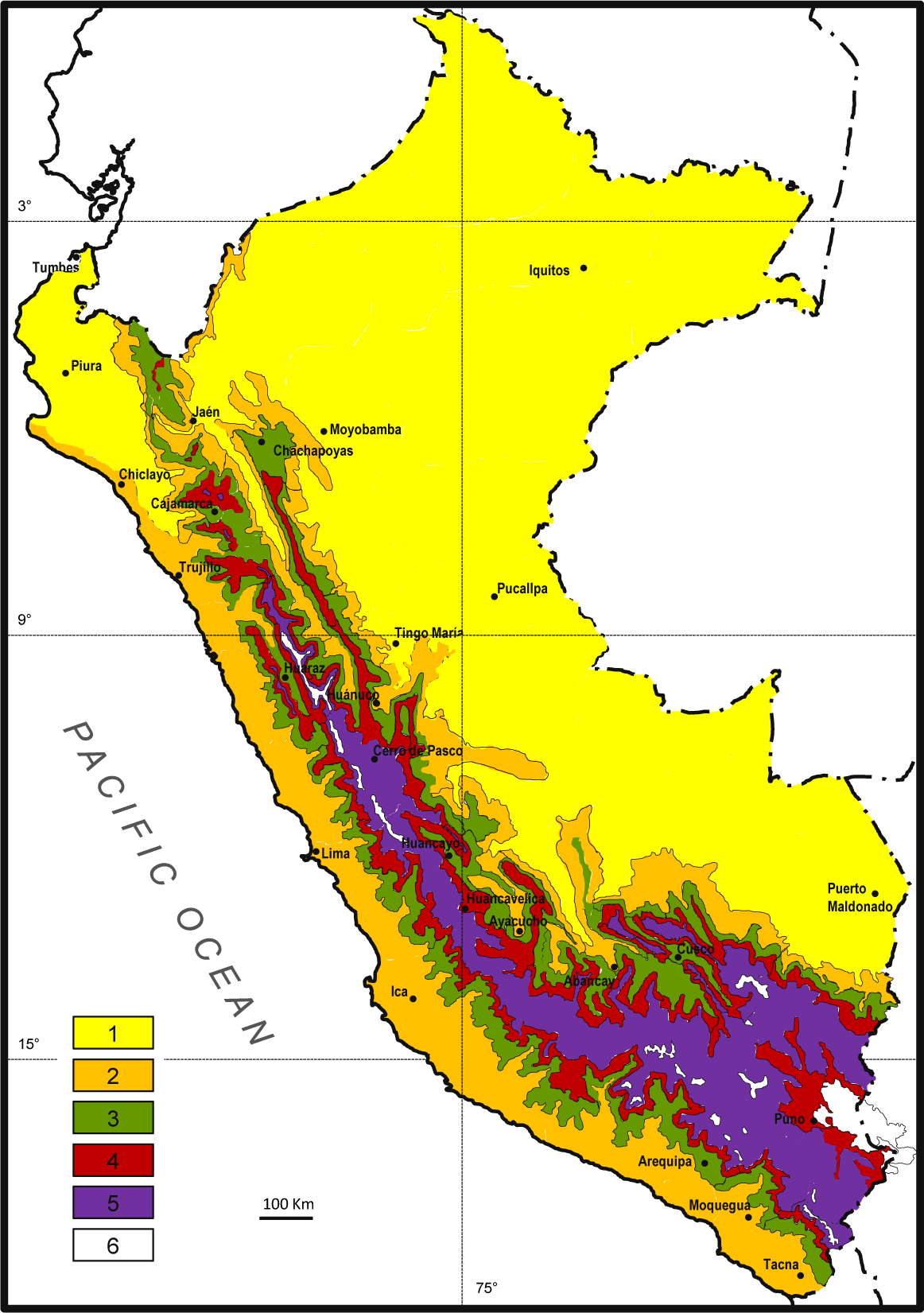
Bioclimatic belts in Peru. 1- Infratropical, 2- Thermotropical, 3- Mesotropical, 4- Supratropical, 5- Orotropical, 6- Cryorotropical.

These belts are nuanced by rainfall intervals (annual P in mm), so that we can observe a very humid infratropical floor in the Amazon but be very dry in northern Peru. We can distinguish nine types of precipitation intervals in the country: Ultrahyperarid (P < 5), hyperarid (5 to 30), arid (31 to 100), semiarid (101 to 300), dry (301 to 500), subhumid (501 to 900), humid (901 to 1500), hyperhumid (1501 to 2500), and ultrahyperhumid (> 2500).

### Biogeographic map of Peru

The bioclimatic belts, with their range of precipitation and plant communities, represent a sequence of vegetation from the base to the summits of the mountains, or from low to high altitudes, also correlated with the nature of the soils and geomorphology, defining an altitudinal cliserie. The cliseries define biogeographic areas (Rivas-Martínez 2007).

The basic biogeography unit is the tessera, which can be defined as a homogeneous territory that can only contain one type of potential vegetation, and therefore, a sequence of substitute communities. It is the only unit that can be repeated disjunctively. The district is a region with peculiar associations that are lacking in nearby districts, where traditional land use is also important. The sector is an area that encompasses several districts and is characterized by its own species and associations. Several sectors make up a biogeographic province, which has its own species, including paleoendemic plants and independent taxa at the genus level; it also contains its own associations and alliances that are organized in a particular altitudinal cliserie. The region is a fairly large territory formed by several biogeographic provinces, so it contains species, genera and even endemic families that form particular phytosociological orders and classes (Rivas-Martínez 1987). Figure 3 shows the altitudinal cliserie of the Arequipa sector within the Xerophytic Punenian province.

**Figure 3.**
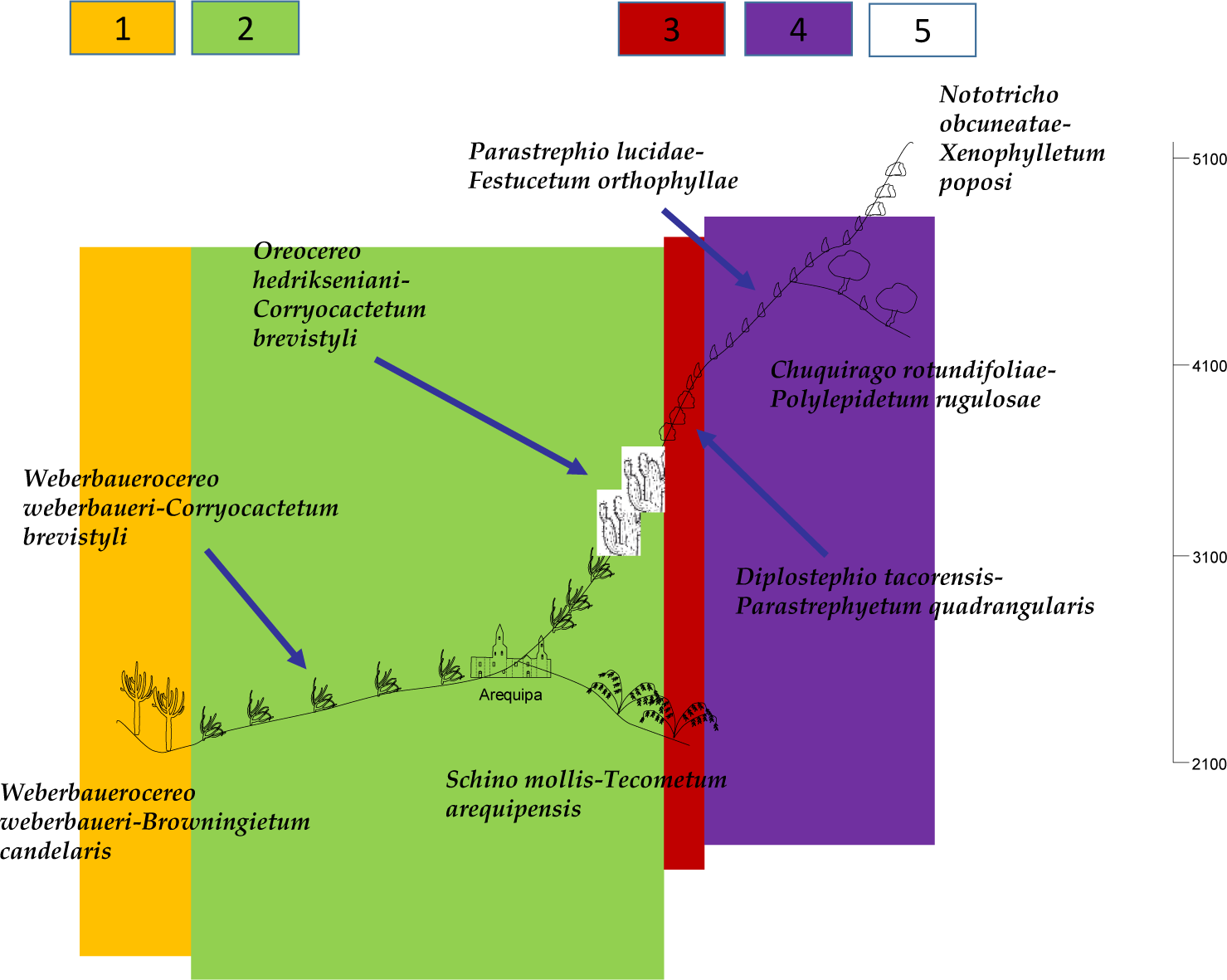
Altitudinal cliserie in the Arequipa sector within the Xerophytic Punenian province. 1- Thermotropical belt, 2- Mesotropical, 3- Supratropical, 4- Orotropical, 5- Cryorotropical.

Our biogeographic division is based on that of Rivas-Martínez et al. (2011) but with certain modifications derived from our experience on bioclimatology, geomorphology, flora and vegetation of Peru (Fig. 4):

**Figure 4.**
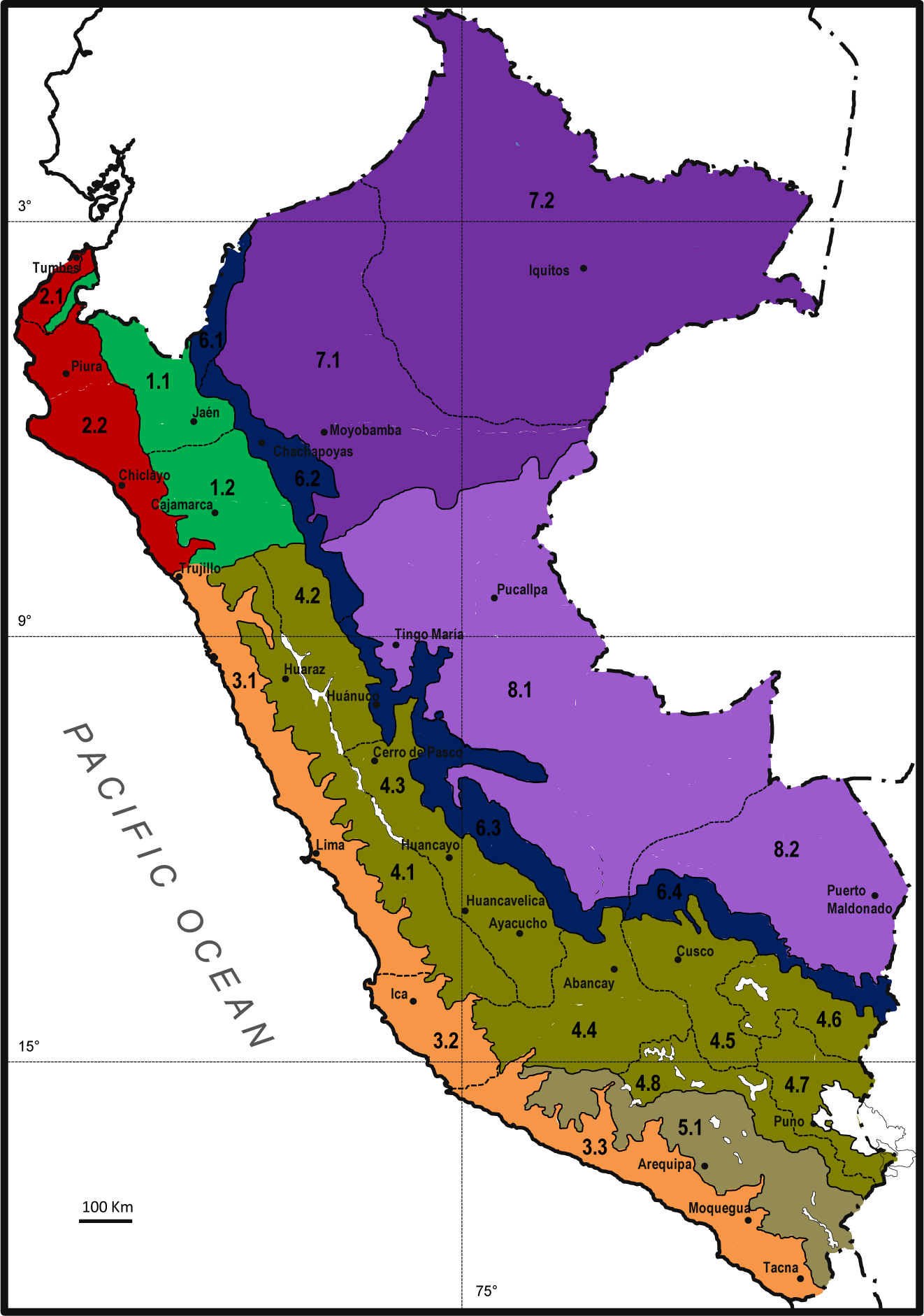
Biogeographic map of Peru. Numbers correspond to those of the biogeographic division in the text.

## NEOTROPICAL-AUSTROAMERICAN KINGDOM

Neotropical Subkingdom

Neogranadian Region

**1. Guayaquilean-Ecuadorean Province:** 1.1. Loja-Cutervo Sector, 1.2. Chota-Contumazá Sector

**2. Pacific Province:** 2.1. Tumbes Sector, 2.2. Sechura Sector

Hyperdesertic Tropical Pacific Region

**3. Hyperdesertic North Peruvian Province:** 3.1. Lima Sector, 3.2. Ica Sector, 3.3. Moquegua-Tacna Sector

Tropical South Andean Region

**4. Mesophytic Punenian Province:** 4.1. Huaraz Sector, 4.2. Chavin Sector, 4.3. Xauxa Sector, 4.4. Apurimac Sector, 4.5. Cusco Sector, 4.6. Allincapac Sector, 4.7. Tititaca Basin Sector, 4.8. Colca Sector.

**5. Xerophytic Punenian Province: Arequipa Sector**

**6. Yungenian Province:** 6.1. North of Peru Sector, 6.2. Chachapoyas Sector, 6.3. Huánuco-Junín Sector, 6.4. Urubamba Sector.

Amazonian Region

**7. West Amazonian Province:** 7.1. Pastaza-Marañón Sector, 7.2. Iquitos Sector

**8. Southwest Amazonian Province:** 8.1. Contamana-Pucallpa Sector, 8.2. Madre de Dios Sector

### Arranging the vegetation classification of Peru

The syntaxonomic scheme of the vegetation of Peru is organized according to the following plant formations: Forests (dry forest, high Andean forest, Andean and Amazonian rainforest), shrublands (Andean shrubland, shrubland associated with watercourses), vegetation with cacti from central and southern Peru, grasslands and scrub of the highlands, vegetation of cryoturbed soils, vegetation of salt marshes (coastal saline soil vegetation, vegetation of inland saline soils), vegetation of maritime dunes, herbaceous vegetation of the coastal hills of the Pacific coast, short annual pioneer vegetation, wetland vegetation (vegetation of depressions, springs and peat bogs; helophyte vegetation, amphibious high Andean vegetation of small plants, amphibious vegetation of small plants from warm areas; floating, temporally rooted, and natant leafy plant communities), rocky and wall vegetation, vegetation of anthropic environments (ruderal and adapted to trampling Andean vegetation, ruderal and adapted to trampling vegetation from the Amazonia and irrigated areas, vegetation of disturbed soils and dumps), and savannas. Within these plant formations, syntaxa are arranged from classes to associations. If syntaxa contain synonyms or other occurrences with the names according to the International Code of Phytosociological Nomenclature, these are included next to the valid name in square brackets, followed by the abbreviations: **Art.:** The Code article by which a name registers an occurrence that invalidates it; **Incl.:** A syntaxon includes another syntaxon, and the one that is included should be combined as a subassociation; **Ined.:** Syntaxon unpublished under study; **Nom. amb.:** *nomen ambiguum*; **Nom. corrigend.:** *nomen corrigendum;* **Nom. dub.:** *nomen dubium;* **Nom. inval.:** *nomen invalidum;* **Nom. invers.:** *nomen inversum;* **Nom. mut. nov.:** *nomen mutatum novum;* **Nom. prov.:** *nomen provisorium;* **Nom. superfl.:** *nomen superfluum;* **p.p.:** A name is a synonym but only in part; **Ref.:** Reference to a text where a syntaxon is found that, due to its floristic composition, must be in Peru, although we have not studied it. **Ref. Amaz.:** References to Amazonian ecological nomenclature; **Syntax. syn.:** Refers to a syntaxon with similar floristic composition.

For each association, a brief ecological description, its bioclimatic affinity, and the biogeographic sector where it was observed according to the numbering of the map in figure 4, are given. The complete bibliography of the syntaxa authorship is in the references section.

### Plant nomenclature

The names of the taxa used were checked using IPNI (2021), The Plant List (2013), and TROPICOS (2021). TROPICOS was also used for the mosses.

### Syntaxonomical scheme for the vegetation of Peru

#### A. FORESTS

##### A.1. DRY FORESTS

###### CLASS I. ACACIO MACRACANTHAE-PROSOPIDETEA PALLIDAE

Galán 1999

+ Acacio macracanthae-Prosopidetalia pallidae Galán 1999

* Tecomion arequipensis Galán & Cáceres in Galán, Rosa & Cáceres 2002

**1. *Acacio macracanthae-Tecometum guarumis*** Galán, Linares, Campos & Vicente 2009

Association of phreatophytes of the streams of the Ica peninsula. Thermo-mesotropical hyperarid. Biogeography: 3.2.

**2. *Schino mollis-Acacietum macracanthae*** Galán & Cáceres in Galán, Rosa & Cáceres 2002

Association of phreatophytes of the rivers that flow down from the Andes to the Peruvian desert. Thermotropical hyperarid. Biogeography: 3.1, 3.2.

* Tecomion fulvae Galán & Cáceres in Galán 1999

**3. *Schino mollis-Prosopidetum calderensis*** ass. nov. Galán & Linares 2020, ined.

Phreatophyte association of areas with gypsiferous outcrops associated with the Caldera Batholith, in Arequipa. Thermo-mesotropical hyperarid-arid. Biogeography: 5.1.

**4. *Schino mollis-Tecometum arequipensis*** Galán, Linares, Campos & Vicente 2009

Association of phreatophytes in areas with sandy soils and volcanic pumices in the surroundings of Arequipa. Thermo-mesotropical hyperarid-arid. Biogeography: 5.1.

**5. *Schino mollis-Tecometum tanaceiiflorae*** Galán, Linares, Campos & Vicente 2009

Association of phreatophytes of the tributary streams of the Colca River in the Department of Arequipa. Thermo-mesotropical hyperarid-arid. Biogeography: 3.3, 5.1.

**6. *Tecometum fulvae Galán*** 1996

Association of phreatophytes of the streams between the Department of Moquegua and northern Chile. Thermo-mesotropical hyperarid. Biogeography: 3.3, 5.1.

+ Cryptocarpo pyriformis-Prosopidetalia pallidae Galán & Cáceres in Galán, Rosa & Cáceres 2002

* Baccharito oblongifoliae-Jacarandion acutifoliae Galán, Sánchez, Linares, Campos, Montoya & Vicente 2016

**7. *Armatocereo balsasensis-Cercidietum praecocis*** Galán, Sánchez, Linares, Campos, Montoya & Vicente 2016

Association of inter-Andean trees and cacti of northern Peru. Infratropical semiarid. Biogeography: 1.1, 1.2.

**8. *Diplopterydo leiocarpae-Acacietum macracanthae*** Galán, Sánchez, Linares, Campos, Montoya & Vicente 2016

Inter-Andean dry forests of northern Peru. Thermotropical dry. Biogeography: 1.1, 1.2.

* Bursero graveolentis-Prosopidion pallidae Galán & Cáceres in Galán, Rosa & Cáceres 2002

**9. *Cercidio praecocis-Prosopidetum pallidae*** Galán & Cáceres in Galán, Rosa & Cáceres 2002

Climatic dry forests of northern Peru. Infra-thermotropical arid-semiarid. Biogeography: 2.1, 2.2.

**10. *Crotono ruiziani-Acacietum macracanthae*** Galán, Sánchez, Linares, Campos, Montoya & Vicente 2016

Western Andean dry forests of northern Peru. Thermotropical dry. Biogeography: 1.2.

**11. *Loxoterygio huasanginis-Neoraimondietum arequipensis*** Galán, Sánchez, Linares, Campos, Montoya & Vicente 2016

Western Andean tree and cactus communities of northern Peru. Infratropical arid-semiarid. Biogeography: 1.2, 2.2.

###### CLASS II. CARICO CANDICANTIS-CAESALPINIETEA SPINOSAE

Galán, Linares, Campos & Vicente 2009

+ Citharexylo flexuosi-Crotonetalia alnifolii Galán, Linares, Campos & Vicente 2009

* Grindelion glutinosae Galán, Linares, Campos & Vicente 2009

**12. *Caesalpinio spinosae-Myrcianthetum ferreyrae*** Galán, Linares, Campos & Vicente 2009 Climatic forests of the coastal hills of southern Peru. Thermotropical dry. Biogeography: 3.3.

**13. *Echinopsio chalaensis-Randietum armatae*** Galán, Linares, Campos & Vicente 2009

Shrub vegetation with cacti in the coastal hills of southern Peru. Thermotropical semiarid. Biogeography: 3.3.

##### A.2. HIGH ANDEAN FORESTS

###### CLASS III. POLYLEPIDETEA TARAPACANO-BESSERI

Rivas-Martínez & Navarro in Navarro & Maldonado 2002

+ Polylepidetalia racemosae Galán & Cáceres in Galán, Rosa & Cáceres 2002 [*Polylepidetalia tarapacano-besseri* Navarro in Navarro & Maldonado 2002, syntax. syn.]

* Polylepidion incano-besseri Navarro in Navarro & Maldonado 2002 [*Polylepidion acuminatae* Galán, Rosa & Cáceres 2002, nom. inval. Art. 3b (prov.)]

**14. *Ageratino azangaroensis-Polylepidetum incarum*** ass. nov. [*Lupino chlorolepis-Polylepidetum incarum* Montesinos-Tubée, Pinto, Beltrán & Galiano 2015b, nom. inval. Art. 4 & 5]

Forest dominated by *Polylepis incarum* on rocky and stony areas from the Titicaca Basin. Supra-orotropical, subhumid. Biogeography: 4.7. Characteristic plants: *Ageratina azangaroensis, Lupinus chlorolepis, Polylepis incarum, Siphocampylus tupaeformis, Stevia macbridei, Viguiera pazensis.* Holotypus: Montesinos-Tubée, Pinto, Beltrán & Galiano 2015b: Tab. 3, plot 5.

**15. *Barnadesio dombeyanae-Polylepidetum racemosae*** ass. nov.

Forest dominated by *Polylepis racemose* on rocky and stony areas of northern Peru (Cajamarca). Supratropical subhumid-humid. Biogeography: 1.2. Characteristic plants: *Barnadesia dombeyana, Buddleja incana, Calceolaria ballotifolia, Polylepis racemosa.* Holotypus: *Barnadesia dombeyana* 1, *Polylepis racemosa* 4, *Buddleja incana* 2, *Calceolaria ballotifolia* 1, *Ageratina sternbergiana* 2, *Baccharis latifolia* 2, *Minthostachys mollis* 1, *Bidens triplinervia* +, *Junellia occulta* 1, *Solanum zahlbruckneri* 1 (Elevation 3595 m, area 100 m^2^, Cajamarca, Nicuipampa, 17M 805409-9221698).

**16. *Mutisio cochabambensis-Polylepidetum incarum*** ass. nov. Galán & Linares 2020, ined.

Forest dominated by *Polylepis incarum* on rocky and stony areas. Supratropical humid. Biogeography: 6.4.

* Ribesido brachybotrys-Polylepidion rugulosae Galán & Cáceres in Galán, Rosa & Cáceres 2002, nom. corrigend. Montesinos-Tubée, Pinto, Beltrán & Galiano 2015b [*Ribesido brachybotrys-Polylepidion besseri*]

**17. *Chuquirago rotundifoliae-Polylepidetum rugulosae*** (Galán, Cáceres & González 2003) Luebert & Gajardo 2005 [*Diplostephio tacorensis-Parastrephietum lepidophyllae polylepidetosum besseri* Galán, Cáceres & González 2003, nom. corrigend. Luebert & Gajardo 2005]

Forest dominated by *Polylepis rugulosa* on rocky and stony areas from the dry Andes of southern Peru. Supra-orotropical, semiarid-dry. Biogeography: 5.1.

* Polylepidion tomentello-tarapacanae Navarro in Navarro & Maldonado 2002

**18. *Mutisio lanigerae-Polylepidetum tarapacanae*** Navarro in Navarro & Maldonado 2002 [*Parastrephio lucidae-Festucetum orthophyllae polylepidetosum tarapacanae* Galán, Cáceres & González 2003, syntax. syn.]

Bushes and grasslands with *Polylepis tarapacana* on rocky and stony areas from the dry Andes of southern Peru. Orotropical semiarid-dry. Biogeography: 5.1.

##### A.3. ANDEAN AND AMAZONIAN RAINFORESTS

Insertae Sedis:

**19. *Oenocarpo maporae-Mauritietum flexuosae*** Galán 1996

Amazon palm grove installed in permanently flooded depressions. Infratropical ultra-hyperhumid. Biogeography: 7.2.

**20. *Pachiro brevipedis-Euterpetum catingae*** Galán 2001

Edapho-hygrophilous palm grove on sandy soils of the lower Amazonia. Infratropical ultra-hyperhumid. Biogeography: 7.2.

###### CLASS IV. ALNETEA ACUMINATAE

Galán 2005

+ Alnetalia acuminatae Galán & Rosa in Galán, Rosa & Cáceres 2002

* Cyatheo herzogii-Alnion acuminatae Galán, Campos, Linares, Montoya, Torres & Vicente 2020

**21. *Hypolepido parallelogrammae-Alnetum acuminatae*** Galán, Campos, Linares, Montoya, Torres & Vicente 2020

Northern Peruvian forest with alders, tree ferns, on black hydromorphic soils with steep slopes. Thermotropical humid. Biogeography: 6.2.

* Escallonio pendulae-Alnion acuminatae Galán, Campos, Linares, Montoya, Torres & Vicente 2020

**22. *Axinaeo tomentosae-Ceroxylonetum quindiuensis*** Galán, Campos, Linares, Montoya, Torres & Vicente 2020

Palm grove dominated by *Ceroxylon quindiuense,* sited on peaty soils. Thermotropical subhumid. Biogeography: 1.1, 6.2.

**23. *Lueheo divaricatae-Ceroxylonetum peruviani*** Galán, Campos, Linares, Montoya, Torres & Vicente 2020

Forest of the endemic palm *Ceroxylum peruvianum* of the Amazonas Department, sited on hydromorphic soils near watercourses and slopes (20–40%) with permanent water. Mesotropical humid. Biogeography: 6.2.

**24. *Mutisio wurdackii-Alnetum acuminatae*** Galán, Campos, Linares, Montoya, Torres & Vicente 2020

Alder forest in northern Peru, found in permanent water courses forming a gallery of trees. Thermotropical subhumid. Biogeography: 6.2.

* Myrico pubescentis-Alnion acuminatae Galán & Rosa in Galán, Rosa & Cáceres 2002

**25. *Valleo stipularis-Alnetum acuminatae*** Galán & Rosa in Galán, Rosa & Cáceres 2002

Alder forest in central Peru, forming a gallery of trees in permanent water courses. Meso-supratropical dry-subhumid. Biogeography: 1.2, 4.1, 4.2, 4.3.

###### CLASS V. MORELLO PUBESCENTIS-MYRSINETEA CORIARIACEAE

Galán, Campos, Linares, Montoya, Torres & Vicente 2020

+ Saurauio peruvianae-Condaminetalia corymbosae Galán, Campos, Linares, Montoya, Torres & Vicente 2020

* Pteridi creticae-Cyatheion caracasanae Galán, Campos, Linares, Montoya, Torres & Vicente 2020

**26. *Viburno reticulati-Weinmannietum spruceanae*** Galán, Campos, Linares, Montoya, Torres & Vicente 2020

Rainforests from northern Peru (Cajamarca Region), on clayey soils of Tertiary volcanic origin and quaternary sediments with a slope up to 60%. Thermotropical humid. Biogeography: 1.1.

* Sanchezio oblongae-Hedyosmion racemose Galán, Campos, Linares, Montoya, Torres & Vicente 2020

**27. *Cecropio albicantis-Weinmannietum glomeratae*** Galán, Campos, Linares, Montoya, Torres & Vicente 2020

Rainforest from central Peru rich in *Weinmannia* species, located on very clayey soils, reddened by iron oxides, with a relief of steep slopes (40–60%). Mesotropical hyperhumid. Biogeography: 6.3.

**28. *Cedrelo fissilis-Ficetum maximae*** Galán, Campos, Linares, Montoya, Torres & Vicente 2020

Rainforest of central Peru periodically flooded with clayey soils with certain hydromorphism. Thermotropical hyperhumid. Biogeography: 6.3.

**29. *Chelyocarpo ulei-Acacietum loretensis*** Galán, Campos, Linares, Montoya, Torres & Vicente 2020

Amazonian rainforests from the lowest parts of the Peruvian Andes, with a hilly relief. They are settled on very yellowish silty soils. Infratropical ultra-hyperhumid. Biogeography: 8.1.

**30. *Piperi chanchamayani-Cecropietum strigosae*** Galán, Campos, Linares, Montoya, Torres & Vicente 2020

Rainforest with *Cecropia* from central Peru (Chanchamayo area) developing on clay-rich soils, with slopes up to 40%. Upper thermotropical hyperhumid. Biogeography: 6.3.

* Serpocaulo dasypleuronis-Alchorneion latifoliae Galán, Campos, Linares, Montoya, Torres & Vicente 2020

**31. *Acalypho macrostachyae-Cecropietum polystachyae*** Galán, Campos, Linares, Montoya, Torres & Vicente 2020

Forest from southern Peru with *Cecropia polystachya* located in flat areas, but also on slopes up to 70% on soils rich in clay. Upper thermotropical ultra-hyperhumid. Biogeography: 6.4.

**32. *Iriartello setigerae-Cinchonetum micranthae*** Galán, Campos, Linares, Montoya, Torres & Vicente 2020

Forest from southern Peru very rich in Peruvian bark trees, located in soft reliefs on clayey soils. Lower thermotropical ultra-hyperhumid. Biogeography: 6.4.

###### CLASS VI. PRUNO RIGIDAE-OREOPANACETEA FLORIBUNDAE

Galán 2005

+ Cestro auriculati-Prunetalia rigidae Galán & Rosa in Galán, Rosa & Cáceres 2002

* Monnino pilosae-Myrcianthion myrsinoidis Galán, Sánchez, Montoya, Linares, Campos & Vicente 2015 [*Symploco nanae-Oreopanacion rusbyi* Galán & Rosa in Galán, Rosa & Cáceres 2002, nom. inv. Art. 3b (prov.)]

**33. *Aristeguietio discoloris-Kageneckietum lanceolatae*** Galán, Sánchez, Montoya, Linares, Campos & Vicente 2015

Andean shrubby sclerophyllous forest of northern Peru. Upper Mesotropical dry-subhumid. Biogeography: 1.2.

**34. *Axinaeo nitidae-Podocarpetum oleifolii*** Galán, Sánchez, Montoya, Linares, Campos & Vicente 2015

Andean laurel-like forests of northern Peru. Mesotropical humid-hyperhumid. Biogeography: 1.2.

**35. *Berberido beauverdianae-Myrcianthetum myrsinoidis*** Galán, Sánchez, Montoya, Linares, Campos & Vicente 2015

Andean sclerophyllous forest of northern Peru. Mesotropical, dry-subhumid. Biogeography: 1.2.

**36. *Crotono churumayensis-Hesperomeletum ferrugineae*** Galán, Campos, Linares, Montoya, Torres & Vicente 2020

Andean forests of southern Peru located on stony soils with scant slopes. Mesotropical humid. Biogeography: 6.4.

**37. *Smallantho parvicipitis-Oreopanacetum eriocephali*** Galán, Campos, Linares, Montoya, Torres & Vicente 2020

Andean forests of southern Peru located on shallow sandy soils on steep slopes. Mesotropical hyperhumid. Biogeography: 6.4.

**38. *Verbesino auriculigerae-Siparunetum muricatae*** Galán, Sánchez, Montoya, Linares, Campos & Vicente 2015

Andean laurel-like forests of anthropogenic origin in northern Peru. Mesotropical humid-hyperhumid. Biogeography: 1.2.

* Myrcianthion quinquelobae Galán & Rosa in Galán, Rosa & Cáceres 2002

**39. *Tristerido peruviani-Myrcianthetum quinquelobae*** Galán & Rosa in Galán, Rosa & Cáceres 2002

Western Andean laurel-like forests in central Peru. Meso-supratropical subhumid-humid. Biogeography: 4.1.

#### B. SCHRUBLANDS

##### B.1. ANDEAN SCHRUBLANDS

###### CLASS VII. CLEMATIDO PERUVIANAE-BACCHARITETEA LATIFOLIAE

Galán, Sánchez,

Montoya, Linares, Campos & Vicente 2015 [*Baccharidetea latifoliae* Lauer, Rafiqpoor & Theisen 2001, nom. inval. Art. 5, 8 & 17]

+ Mutisio acuminatae-Baccharitetalia latifoliae Galán & Cáceres in Galán, Rosa & Cáceres 2002

* Mutisio acuminatae-Ophryosporion peruviani Galán & Cáceres in Galán, Rosa & Cáceres 2002

**40. *Aristeguietio discoloris-Baccharitetum latifoliae*** Galán, Baldeón, Beltrán, Benavente & Gómez 2004

Andean shrubs that border the ‘queñual’ trees of central Peru. Supratropical dry-subhumid. Biogeography: 4.1.

**41. *Barnadesio blakeanae-Ophryosporietum peruviani*** Galán & Cáceres in Galán, Rosa & Cáceres 2002

Shrubs of the western Andean slopes of central Peru. Mesotropical dry-subhumid. Biogeography: 4.1.

**42. *Cynancho tarmensis-Tecometum sambucifoliae*** Galán 1996

Shrubs on the boundaries of vegetable gardens in central Peru. Mesotropical dry. Biogeography: 4.3.

**43. *Dunalio spinosae-Baccharitetum latifoliae*** Galán, Cáceres & González 2003

Shrubs of the western valleys of southern Peru. Meso-supratropical dry-subhumid. Biogeography: 4.8, 5.1.

* Otholobio munyensis-Rubion robusti Galán, Sánchez, Montoya, Linares, Campos & Vicente 2015

**44. *Baccharito latifoliae-Monactinetum flaverioidis*** Galán, Sánchez, Montoya, Linares, Campos & Vicente 2015

Heliophilous shrubs bordering alder and ‘queñual’ tree forests of northern Peru. Meso-supratropical dry-subhumid. Biogeography: 1.2.

**45. *Monactino flaverioidis-Colignonietum parviflorae*** Galán, Sánchez, Montoya, Linares, Campos & Vicente 2015

Shrubs bordering the humid forests of northern Peru. Mesotropical humid. Biogeography: 1.2.

##### B.2. SHRUBLANDS ASSOCIATED TO WATERCOURSES

###### CLASS VIII. TESSARIO INTEGRIFOLIAE-BACCHARITETEA SALICIFOLIAE

Rivas-Martínez & Navarro in Navarro & Maldonado 2002

+ Plucheo absinthioidis-Baccharitetalia salicifoliae Rivas-Martínez & Navarro in Navarro & Maldonado 2002 [*Baccharitetalia salicifoliae* Galán, Baldeón, Beltrán, Benavente & Gómez 2004, syntax. syn.]

* Pityrogrammo trifoliatae-Baccharition salicifoliae Galán, Baldeón, Beltrán, Benavente & Gómez 2004 [*Taraso capitatae-Salicion humboldtianae* Montesinos-Tubée & Núñez in Montesinos-Tubée, Núñez, Toni, Álvarez, Borgoño, Zegarra, Gutiérrez, Maldonado, Rodríguez, Riveros & Guillén 2019, syntax. syn., nom. inval. Art. 5, nom. dub. Art. 37]

**46. *Baccharito salicifoliae-Gynerietum sagittati*** Galán, Baldeón, Beltrán, Benavente & Gómez 2004 [*Baccharis salicifolia-Tessaria integrifolia* Ges. Müller & Gutte 1985, nom. inval. Art. 3c]

Shrubs with tall helophytes on stony riverbeds from the western Andes and coastal desert. Termo-mesotropical árido. Biogeografía: 3.1, 3.2, 3.3.

**47. *Equiseto gigantei-Salicetum humboldtianae*** Galán, Baldeón, Beltrán, Benavente & Gómez 2004 [*Taraso capitatae-Baccharidetum salicifoliae* Montesinos-Tubée & Núñez in Montesinos-Tubée, Núñez, Toni, Álvarez, Borgoño, Zegarra, Gutiérrez, Maldonado, Rodríguez, Riveros & Guillén 2019, syntax. syn., nom. inval. Art. 3k, 10b, 5 & 16, nom. dub. Art. 37; *Helogyno stramineae-Muehlenbeckietum hastulatae* Montesinos-Tubée & Núñez in Montesinos-Tubée, Nuñez, Toni, Álvarez, Borgoño, Zegarra, Gutiérrez, Maldonado, Rodríguez, Riveros & Guillén 2019, syntax. syn., nom. inval. Art. 5, nom. dub. Art. 37]

Willow groves on the sandy sediments of the rivers that flow down from the Andes to the Pacific coast. Thermo-mesotropical, hyperarid-arid. Biogeography: 3.1, 3.3, 5.1.

* Plucheion absinthioidis Galán, Linares, Campos & Vicente 2009

**48. *Geoffroeetum decorticantis*** ass. nov. [*Gourlietum* Pisano 1956, nom. inval. Art. 2 & 7; Com.

*Geoffroea decorticans* Galán, Linares, Campos & Vicente 2009, nom. inval. Art. 3c]

Flooded plant communities formed by shrubs living on salty soils that form polygonal crusts when they dry out. Thermotropical hyperarid. Biogeography: 3.3. Characteristic plants: *Baccharis salicifolia, Geoffroea decorticans, Pluchea absinthioides*. Holotypus: *Geoffroea decorticans* 3, *Pluchea absinthioides* +, *Baccharis salicifolia* 2, *Distichlis spicata* 1, *Schinus molle* 1, *Acacia macracantha* 2 (Elevation 200 m, area 100 m^2^, Tacna, quebrada de Sama, 19K 323456-7991230).

**49. *Lycio distichum-Baccharitetum uniflorae*** Galán, Linares, Campos, Trujillo, Villasante & Vicente 2011b

Shrubs on sloping gorges with saline efflorescence. Thermotropical arid. Biogeography: 5.1.

**50. *Plucheetum absinthioidis*** Galán, Linares, Campos & Vicente 2009 Coastal shrubs on saline soils. Thermotropical hyperarid. Biogeography: 3.3.

##### C. VEGETATION WITH CACTI FROM CENTRAL AND SOUTHERN PERU

###### CLASS IX. OPUNTIETEA SPHAERICAE

Galán & Vicente 1996 [*Haageocereetea acranthi* Müller 1985b, nom. inval. Art. 3b (prov.); *Echinopsio schoenii-Proustietea cuneifoliae* Montesinos-Tubée, Cleef & Sýkora 2012, nom. inval. Art. 8 & 9]

+ Oreocereo leucotrichi-Neoraimondietalia arequipensis Galán & Vicente 1996 [*Haageocereetalia olowinskiani* Müller 1985b, nom. inval. Art. 3b (prov.)]

* Corryocaction brevistyli Galán & Vicente 1996

**51. *Armatocereo riomajensis-Neoraimondietum arequipensis*** Galán, Linares, Campos, Trujillo, Villasante & Vicente 2011b

Columnar cactus vegetation on ignimbrites and conglomerates from the inland valleys of the Arequipa Department. Upper thermotropical arid. Biogeography: 4.8.

**52. *Balbisio weberbaueri-Ambrosietum artemisioidis*** Galán, Linares, Campos, Trujillo, Villasante & Vicente 2011b

Scrublands with cacti on the western slopes of the volcanic environment of the Arequipa Department. Upper mesotropical semiarid. Biogeography: 5.1.

**53. *Corryocacto aurei-Browningietum candelaris*** Galán & Vicente 1996

Columnar cactus vegetation of the western Andean slopes of the Tacna Department. Mesotropical hyperarid. Biogeography: 5.1.

**54. *Grindelio bolivianae-Corryocactetum puquiensis*** Galán & Gómez 2001

Columnar cactus vegetation from the inner valleys of the Arequipa Department. Meso-supratropical arid-semiarid. Biogeography: 4.8.

**55. *Neoraimondio arequipensis-Browningietum viridis*** Galán, Linares, Campos, Trujillo, Villasante & Vicente 2011b

Inter-Andean association on limestones and quartzite conglomerates from the inland valleys of the Arequipa Department. Thermotropical arid. Biogeography: 4.8.

**56. *Oreocereo tacnaensis-Corryocactetum brevistyli*** Galán & Vicente 1996

Columnar cactus vegetation from the western Andean slopes of the Tacna Department. Mesotropical arid-semiarid. Biogeography: 5.1.

**57. *Weberbauerocereo rauhii-Browningietum candelaris*** Galán, Linares, Campos & Vicente 2009

Columnar cactus vegetation from the western Andean slopes of the Arequipa Department. Thermotropical arid. Biogeography: 5.1.

**58. *Weberbauerocereo rauhii-Corryocactetum brevistyli*** Galán, Linares, Campos & Vicente 2009

Columnar cactus vegetation from the western Andean slopes of the Arequipa Department. Mesotropical arid-semiarid. Biogeography: 5.1.

**59. *Weberbauerocereo torataensis-Corryocactetum brevistyli*** Galán, Linares, Campos & Vicente 2009

Columnar cactus vegetation from the western Andean slopes of the Torata River basin (Moquegua). Mesotropical arid. Biogeography: 5.1.

**60. *Weberbauerocereo weberbaueri-Browningietum candelaris*** Galán & Linares 2012

Columnar cactus vegetation from the western Andean slopes of the volcanic environment of the Arequipa city. Thermotropical arid. Biogeography: 5.1.

**61. *Weberbauerocereo weberbaueri-Corryocactetum brevistyli*** Galán & Gómez 2001

Mesotropical columnar cactus vegetation from the Arequipa Department. Mesotropical arid-semiarid. Biogeography: 5.1.

* Espostoo melanostelis-Neoraimondion arequipensis Galán & Rosa in Galán, Rosa & Cáceres 2002

**62. *Haageocereo limensis-Neoraimondietum arequipensis*** Galán & Rosa in Galán, Rosa & Cáceres 2002

Columnar cactus vegetation from the western Andean slops of Central Peru. Thermotropical arid. Biogeography: 4.1.

**63. *Haageocereo versicoloris-Armatocereetum proceri*** Galán, Sánchez, Linares, Campos, Montoya & Vicente 2016

Association of columnar cacti in areas with maritime influence in central Peru. Thermotropical hyperarid. Biogeography: 3.1.

* Haageocerion australis Galán, Cáceres & González 2002

**64. *Neoporterio islayensis-Neoraimondietum arequipensis*** Galán, Cáceres & González 2002

Columnar cactus communities of the southern Peruvian desert. Thermotropical hyperarid. Biogeography: 3.3.

* Haageocerion acranthi all. nov. [*Haageocerion olowinskiani* Müller 1985b, nom. inval. Art. 3b (prov.)]

Alliance that gathers associations of Cactaceae rich in geophytes and lichens by condensation of fogs in more or less stony inland areas of the hills of the coastal desert of central Peru. Characteristic plants: *Alstroemeria pelegrina, Anredera diffusa, Atriplex espostoi, Echeandia eccremorrhiza, Haageocereus acranthus, Loxanthocereus acanthurus, Oxalis megalorrhiza.* Holotypus: *Echeandio eccremorrhizae-Haageocereetum acranthi* Müller 1985b.

**65. *Echeandio eccremorrhizae-Haageocereetum acranthi*** Müller 1985b nom. invers. Art. 42, nom. mut. nov. [*Haageocereo olowinskiani-Anthericetum eccremorrhizi*]

Cactus communities of the central region of the Peruvian desert. Thermotropical hyperarid. Biogeography: 3.1.

+ Echinopsio schoenii-Proustetalia cuneifoliae ord. nov. [*Echinopsio schoenii-Proustetalia couneifoliae* Montesinos-Tubée, Cleef & Sýkora 2012, nom. inval. Art. 8 & 9]

This order includes meso-supropical associations with columnar cacti and scrublands in the inland valleys of Arequipa and Moquegua. Characteristic plants: *Cylidropuntia imbricata* subsp. *rosea, Echinopsis schoenii, Euphorbia apurimacensis, Junellia arequipense, Proustia cuneifolia, Senecio arnaldii.* Holotypus: *Echinopsio schoenii-Proustion cuneifoliae* all. nov.

* Echinopsio schoenii-Proustion cuneifoliae all. nov. [*Salvion oppositiflorae* Montesinos-Tubée, Cleef & Sýkora 2012, nom. inval. Art. 38]

Holotypus: *Junellio arequipensis-Proustietum cuneifolii* ass. nov.

**66. *Anredero diffusae-Diplostephietum meyenii*** Montesinos-Tubée, Cleef & Sýkora 2012

Association with cacti and shrubs on slopes and flat areas on loose volcanic substrates, in the inland valleys of Moquegua. Supratropical semiarid. Biogeography: 5.1.

**67. *Armatocereo riomajensis-Euphorbietum apurimacensis*** Galán, Linares, Campos & Vicente 2009

Vegetation of columnar cacti and spurge shrubs from the inland valleys of the Arequipa Department. Mesotropical semiarid. Biogeography: 4.8.

**68. *Junellio arequipensis-Proustietum cuneifoliae*** ass. nov. [*Senecioni arnaldii-Exhalimolobetum weddellii* Montesinos-Tubée, Cleef & Sýkora 2012, nom. inval. Art. 3k, nom. dub. Art. 37; incl. *Mostacillastro gracile-Chuquiragetum rotundifoliae* Montesinos-Tubée, Cleef & Sýkora 2012, nom. dub. Art. 37]

Vegetation of columnar cacti and scrubs on steep slopes (up to 50%) with volcanic substrates, in the inland valleys of Arequipa and Moquegua. Supratropical dry. Biogeography: 5.1. Characteristic plants: *Echinopsis pampana, E. schoenii, Gochnatia arequipensis, Junellia arequipense, Neowerdermannia peruviana, Proustia berberidifolia, P. cuneifolia, Senecio arnaldii.* Holotypus: Montesinos-Tubée, Cleef & Sýkora 2012: Tab. 4, plot. 2.

##### D. GRASSLANDS AND SCRUBS OF THE HIGHLANDS

###### CLASS X. CALAMAGROSTIETEA VICUNARUM

Rivas-Martínez & Tovar 1982 [*Luzulo racemosae-Calamagrostietum vicunarum* Gutte 1981, nom. inval. Art. 1; *Calamagrostietea vicunarum* Gutte 1985, nom. inval. Art. 5, 8 & 17.; *Stipetea mucronatae* Gutte 1986, nom. inval. Art. 5, 8 & 17]

+ Agrostio tolucensis-Paspaletalia bonplandiani Galán, Sánchez, Montoya, Linares, Campos & Vicente 2015

* Agrostio tolucensis-Paspalion bonplandiani Galán, Sánchez, Montoya, Linares, Campos & Vicente 2015

**69. *Agrostio tolucensis-Paspaletum bonplandiani*** Galán, Sánchez, Montoya, Linares, Campos & Vicente 2015

Very humid Páramo grassland with gravelly soils derived from granites, where *Puya fastuosa* is abundant. Orotropical humid-hyperhumid. Biogeography: 1.2.

**70. *Calamagrostio tarmensis-Hypericetum laricifolii*** Galán, Sánchez, Montoya, Linares, Campos & Vicente 2015

Grassland with *Hypericum* settled on andosols derived from granite in the Páramo of northern Peru. Supratropical humid. Biogeography: 1.2.

**71. *Oreobolo goeppingieri-Hypericetum laricifolii*** Sabogal [2014] ex Galán, Sánchez, Montoya, Linares, Campos & Vicente 2015

Grassland with Ericaceae settled on andosols derived from granite in the Páramo of northern Peru. Supratropical humid. Biogeography: 1.1.

+ Calamagrostietalia vicunarum Rivas-Martínez & Tovar 1982 [*Calamagrostietalia vicunarum*

Gutte 1985, nom. inval. Art. 5, 8 & 17, syntax. syn.; *Stipetalia ichu* Gutte 1986, nom. inval. Art. 5, 8 & 17]

* Calamagrostion rigidae Gutte 1985, nom. mut. nov. [*Calamagrostion antonianae*] [lectotypus hoc loco: *Festuco dolichophyllae-Calamagrostietum antonianae* Gutte 1985]

**72. *Calamagrostio amoenae-Poetum gymnanthae*** Gutte 1985

Grassland with ‘tola’ (*Parastrephia quadrangularis*) in dry areas exposed to the wind. Orotropical dry. Biogeography: 4.3.

**73. *Festuco dolichophyllae-Calamagrostietum rigidae*** Gutte 1985, nom. mut. nov. [*Festuco dolichophyllae-Calamagrostietum antonianae*]

Tall grasslands of the humid Puna that develop on deep limestone soils, volcanic materials, and sandstones. Orotropical humid. Biogeography: 4.1, 4.3.

**74. *Alchemillo pinnatae-Stipetum hans-meyeri*** ass. nov. [*Lachemillo cf. andinae-Stipetum hans-meyeri* Gutte 1985, nom. inval. Art. 3g]

Grasslands on gravelly soils with steep slopes, solidified on andesitic lava flows. Orotropical humid. Biogeography: 4.3. Characteristic plants: *Alchemilla pinnata, Baccharis ganistelloides, Lupinus conicus, Muehlenbeckia volcanica, Senecio modestus, Stipa hans-meyeri, Valeriana erikae.* Holotypus: Gutte 1985: Tab. 7, plot 4.

**75. *Orthrosantho occissapungi-Puyetum raimondii*** Gutte 1985

Association of very stony slopes and eroded soils of the humid Puna. Orotropical humid. Biogeography: 4.3, 4.7.

* Calamagrostion minimae Rivas-Martínez & Tovar 1982 [*Calamagrostion vicunarum* Gutte 1985, nom. inval. Art. 5, 8 & 17]

**76. *Aciachno pulvinatae-Calamagrostietum vicunarum*** Gutte 1985 [*Gnaphalio badii-Aciachnetum pulvinatae* Seibert 1993, syntax. syn.; *Stipa hans-meyeri-Dissanthelium peruvianum* Ges. Gutte 1985, nom. inval. Art. 3c; *Pycnophyllum aff. lechlerianum* Ges. Gutte 1985, nom. inval. Art. 3c]

Frequently grazed pastures. Orotropical subhumid-humid. Biogeography: 4.1, 4.3.

**77. *Azorello diapensioidis-Calamagrostietum minimae*** Rivas-Martínez & Tovar 1982 [*Baccharito caespitosae-Azorelletum diapensioidis* Montesinos-Tubée, Cleef & Sýkora 2015a, nom. dub. Art. 37; *Astragalo minimi-Azorelletum diapensioidis* Montesinos-Tubée, Cleef & Sýkora 2021, syntax. syn.]

Grassland of polygonal clayey, humid, and deep soils. Oro-cryorotropical subhumid-humid. Biogeography: 4.1, 4.3, 4.8.

**78. *Calamagrostio minimae-Trichophoretum rigidae*** Galán, Méndez, Linares, Campos & Vicente 2014

Association of stony soils through which runoff water of snowmelt flows. Cryorotropical humid. Biogeography: 4.6, 4.7.

**79. *Nototricho pinnatae-Lachemilletum frigidae*** Galán, Méndez, Linares, Campos & Vicente 2014

Grasslands on rock ledges and polygonal soils of central Peru. Cryorotropical subhumid-humid. Biogeography: 4.1.

**80. *Paspalo tuberosi-Piptochaetietum featherstonei*** Gutte 1985, nom. mut. nov. [*Paspalo tuberosi-Piptochaetietum juninensis*]

Grassland on loamy soil derived from Mesozoic limestones. Supra-orotropical dry-subhumid. Biogeography: 4.3.

81. *Pycnophyllo mollis-Festucetum rigescentis* Seibert 1993

Association of flat or sloping areas with solifluction terracing. Cryorotropical subhumid-humid. Biogeography: 5.1.

* Festucion distichovaginatae Gutte 1986 [lectotypus hoc loco: *Muhlenbergio angustatae-Festucetum distichovaginatae* Gutte 1986; *Festucion dichocladae* Gutte 1986, syntax. syn.]

**82. *Muhlenbergio angustatae-Festucetum distichovaginatae*** Gutte 1986

Association of rendzinas, sometimes on steep slopes, of central Peru. Supratropical dry-subhumid. Biogeography: 4.3.

**83. *Oroyo peruvianae-Festucetum distichovaginatae*** Gutte 1986, nom. mut. nov. [*Oroyo suboccultae-Festucetum distichovaginatae*]

Association of terraces, with loose soil, formed by the erosion of limestone materials. Supratropical dry-subhumid. Biogeography: 4.3.

**84. *Elymo cordillerani-Calamagrostietum tarmensis*** Gutte 1986, nom. mut. nov. [*Agropyro attenuati-Calamagrostietum tarmensis*]

Ombrophilous association of shallow limestone soils with preferential South West orientation. Supratropical dry-subhumid. Biogeography 4.3.

**85. *Elymo angulati-Festucetum dichocladae*** Gutte 1986

Heliophilous and grazed association of shallow limestone soils with preferential exposition to North. Supratropical dry-subhumid. Biogeography: 4.3.

+ Coreopsietalia fasciculatae ord. nov. [Sin: *Coriopsietalia fasciculatae* Gutte 1986, nom. inval. Art. 5, 8, 9 & 17]

Grasslands on basic substrates dominated by *Plantago extensa*, on both limestone and Tertiary conglomerates. Characteristic plants: *Castilleja virgatoides, Coreopsis fasciculate, Festuca humilior, Gentianella cuspidate, Plantago extensa, Quinchamalium chilense.* Holotypus: *Plantaginion extensae* Gutte 1986.

* Plantaginion lanuginosae Gutte 1986, nom. mut. nov. [*Plantaginion extensae*] [lectotypus hoc loco: *Altensteinio mathewsii-Plantaginetum extensae* Gutte 1986]

**86. *Ao mathewsii-Plantaginetum lanuginosae*** Gutte 1986, nom. mut. nov. [*Altensteinio mathewsii-Plantaginetum extensae*]

Grasslands on well-formed limestone soils, which occupy large areas in central Peru. Supratropical subhumid-humid. Biogeography: 4.3.

**87. *Astragalo dombeyi-Plantaginetum lanuginosae*** Gutte 1986, nom. mut. nov. [*Astragalo dombeyi-Plantaginetum extensae*]

Grasslands on limestone conglomerates, very localized in central Peru. Supratropical subhumid-humid. Biogeography: 4.3.

* Argyrochosmo niveae-Neowerdermannion peruvianae Montesinos-Tubée, Cleef & Sýkora 2015a

**88. *Plantagini sericeae-Plazietum daphnoidis*** Montesinos-Tubée, Cleef & Sýkora 2015a Scrublands with grasses and rock plants on steep slopes. Supratropical dry. Biogeography: 4.8.

**89. *Neowerdermannio peruvianae-Festucetum weberbaueri*** Montesinos-Tubée, Cleef & Sýkora 2015a

Scrublands with grasses and rock plants on steep slopes. Lower Orotropical dry. Biogeography: 4.8.

+ Parastrephietalia quadrangularis Navarro 1993, nom. mut. Galán, Linares, Campos, Trujillo, Villasante & Vicente 2011b [*Parastrephietalia lepidophyllae*]

* Fabianion stephanii Galán, Cáceres & González 2003

**90. *Diplostephio tacorensis-Parastrephietum quadrangularis*** Galán, Cáceres & González 2003, nom. mut. nov. [*Diplostephio tacorensis-Parastrephietum lepidophyllae*]

Scrub that sits on stony soils with small pebbles from plains and ancient volcanic lava flows. Supratropical semiarid-arid. Biogeography: 5.1.

* Azorello compactae-Festucion orthophyllae Galán, Cáceres & González 2003

**91. *Astragalo pusilli-Parastrephietum quadrangularis*** Montesinos-Tubée, Cleef & Sýkora 2021

*Parastrephia* scrub with some herbs whose presence is favored by intensive grazing. Orotropical dry-subhumid. Biogeography: 4.8, 5.1.

**92. *Baccharito tricuneatae-Puyetum raimondii*** Galán, Linares, Campos & Vicente 2009 Association of very stony andesitic slopes. Orotropical subhumid. Biogeography: 4.8.

**93. *Calamagrostio trichophyllae-Azorelletum compactae*** Montesinos-Tubée, Cleef & Sýkora 2021

Plant community widely distributed on steep rocky slopes with scree and plateaus, where *Calamagrostis trichophylla* replaces *Festuca orthophylla*. Upper orotropical dry-subhumid. Biogeography: 4.8, 5.1.

**94. *Diplostephio tovarii-Festucetum orthophyllae*** Galán, Linares, Campos & Vicente 2009 Inland grassland with scrubs on volcanic soils. Orotropical dry-subhumid. Biogeography: 4.8.

**95. *Parastrephio lucidae-Festucetum orthophyllae*** Galán, Cáceres & González 2003

Western grassland with scrubs on scree and ashes of volcanic origin. Oro-cryorotropical dry. Biogeography: 5.1.

**96. *Parastrephio quadrangularis-Festucetum dolichophyllae*** Galán, Linares, Campos, Trujillo, Villasante & Vicente 2011b [*Sisyrinchio rigidifolii-Parastrephietum quadrangularis* Montesinos-Tubée, Cleef & Sýkora 2015a, syntax. syn.; *Paronychio ubinensis-Hypochaeridetum mucidae* Montesinos-Tubée, Cleef & Sýkora 2015a, nom. inval. Art. 3k; nom. dub. Art. 37; *Festuco dolichophyllae-Nordenstamietum longistylae* Chicalla 2017, nom. inval. Art. 3k, 5 & 10b; incl. *Plantago sericeae-Gnaphalietum sp.* Chicalla 2017, nom. inval. Art. 3b (prov.), 3g]

Grassland with *Parastrephia* scrubs on tuffaceous sandstones and alluvial deposits. Lower Orotropical subhumid. Biogeography: 4.3, 4.4, 4.5, 4.7, 4.8.

**97. *Senecioni moqueguensis-Pycnophylletum mollis*** Montesinos-Tubée, Cleef & Sýkora 2021

Vegetation characterized by the endemic *Senecio moqueguensis,* with cushion plants on bare soils formed by very small andesitic stones. Upper orotropical dry-subhumid. Biogeography: 4.8, 5.1.

**98. *Senecioni nutantis-Parastrephietum quadrangularis*** Galán, Linares, Campos, Trujillo, Villasante & Vicente 2011b

Western scrubland settled on lava fronts and Pleistocene alluvial materials that form fixed scree and deep soils. Low Orotropical dry. Biogeography: 5.1.

##### E. VEGETATION OF CRYOTURBED SOILS

###### CLASS XI. ANTHOCHLOO LEPIDULAE-DIELSIOCHLOETEA FLORIBUNDAE

Rivas-Martínez & Tovar 1982 [*Senecionetea adenophylloidis* Gutte 1987a, nom. inval. Art. 3b (prov.); *Pycnophylletea weberbaueri* Gutte 1987a, nom. inval. Art. 3b (prov.)]

+ Anthochloo lepidulae-Dielsiochloetalia floribundae Rivas-Martínez & Tovar 1982

* Nototrichion obcuneatae Galán, Cáceres & González 2003

**99. *Belloo piptolepis-Dissanthelietum calycini*** Galán, Cáceres & González 2003 Association on flat polygonal soils. Cryorotropical subhumid-humid. Biogeography: 5.1.

**100. *Nototricho obcuneatae-Mniodetum coarctatae*** Galán, Méndez, Linares, Campos & Vicente 2014

Association on polygonal, flat soils, rich in small pebbles and volcanic ashes. Cryorotropical dry-subhumid. Biogeography: 5.1.

**101. *Nototricho obcuneatae-Xenophylletum poposi*** Galán, Cáceres & González 2003

Association on semi-fixed screes due to gelifraction processes. Cryorotropical dry-subhumid. Biogeography: 5.1.

**102. *Poo aequiglumae-Xenophylletum dactylophylli*** Montesinos-Tubée, Cleef & Sýkora 2021

Association on screes with semi-fixed blocks from humid summits. Cryorotropical humid. Biogeography: 4.8, 5.1.

* Xenophyllo ciliolati-Englerocarion peruvianae Rivas-Martínez & Tovar 1982, nom. mut. Galán, Méndez, Linares, Campos & Vicente 2014 [*Wernerio ciliolatae-Englerocharion peruvianae*] [*Wernerion decorae* Gutte 1987a, syntax. syn.; *Wernerion dactylophyllae* Gutte 1987a, p.p., nom. inval. Art. 5 & 17; *Drabion argenteae* Gutte 1987a, p.p. nom. inval. Art. 5, 8 & 17; *Gentianellion vaginalis* Gutte 1987a, p.p., nom. inval. Art. 3c, 5, 8 & 17]

**103. *Anthochloo lepidulae-Culcitietum canescentis*** Gutte 1987a

Association on edges of big rocks. Cryorotropical subhumid-humid. Biogeography: 4.1, 4.3.

**104. *Niphogetono scabrae-Leucherietum daucifoliae*** Gutte 1987a

Vegetation on scree slopes strongly exposed to wind and snow. Orotropical subhumid-humid. Biogeography: 4.3.

**105. *Pynophyllo mollis-Valerianetum pycnanthae*** Gutte 1987a

Association on gravel interspersed with clayey materials forming deep soils. Cryorotropical subhumid-humid. Biogeography: 4.3.

**106. *Stangeo rhizanthae-Catadysietum rosulantis*** Rivas-Martínez & Tovar 1982

Association on polygonal soils rich in reddish-brown clays. Cryorotropical subhumid-humid. Biogeography: 4.1, 4.3.

**107. *Xenophyllo ciliolati-Plettkeetum cryptanthae*** Rivas-Martínez & Tovar, nom. mut. Galán, Méndez, Linares, Campos & Vicente 2014 [*Wernerio ciliolatae-Plettkeetum cryptanthae*] [*Wernerietum decorae* Gutte 1987a, syntax. syn.]

Association on scree slopes and moraine deposits of the humid Puna. Cryorotropical subhumid-humid. Biogeography: 4.1, 4.3, 4.6.

#### G. VEGETATION OF SALT MARSHES

##### G.1. COASTAL SALINE SOIL VEGETATION

###### CLASS XII. RHIZOPHORETEA MANGLE

Bolòs, Cervi & Hatschbach 1991

+ Rhizophoretalia mangle Bolòs, Cervi & Hatschbach 1991 [*Rhizophoretalia* Cuatrecasas 1958, nom. inval. Art. 2b & 8]

* Rhizophorion mangle Bolòs, Cervi & Hatschbach 1991 [*Rhizophorion* Cuatrecasas 1958, nom. inval. Art. 2b & 8]

**108. *Pelliciero rhizophorae-Rhizophoretum mangle*** Cortés & Rangel 2011 (association to be confirmed in Peru, ref.: Cuatrecasas 1958, García-Fuentes *et al*. 2020)

Permanently flooded mangroves of northern South America. Infratropical semiarid-arid. Biogeography: 2.1, 2.2.

* Lagunculario racemosae-Avicennion germinantis Peinado, Alcaraz & Delgadillo 1995 [*Rhizophorion* Cuatrecasas 1958, nom. inval. Art. 2b & 8; *Avicennion occidentalis* Cuatrecasas 1958 sensu Borhidi, Muñiz & Del Risco 1979, nom. inval. Art. 2b]

**109. *Batido maritimae-Avicennietum germinantis*** Borhidi & Del Risco in Borhidi 1991 [*Batido-Avicennietum germinantis* Borhidi & Del Risco in Borhidi, Muñiz & Del Risco 1979, nom. inval. Art. 5 & 7; *Batido maritimae-Avicennietum germinantis* Peinado, Alcaraz & Delgadillo 1995, syntax. syn.] (association to be confirmed in Peru, ref.: García-Fuentes et al. 2020)

Temporarily flooded mangroves of Central America, the Caribbean and northern South America. Infratropical semiarid-arid. Biogeography: 2.1, 2.2.

* Conocarpodo-Laguncularion Borhidi 1996 [*Rhizophorion* Cuatrecasas 1958, nom. inval. Art. 2b & 8; *Conocarpo-Laguncularion* Borhidi in Borhidi, Muñiz & Del Risco 1979, nom. inval. Art. 5 & 7;

*Conocarpo-Laguncularion* Borhidi 1991, nom. inval. Art. 5; *Conocarpodo-Laguncularion* Borhidi in García-Fuentes *et al*. 2020, nom. superfl. Art. 18, 29c]

**110. *Lagunculario racemosae-Conocarpodetum erecti*** Peinado, Alcaraz & Delgadillo 1995 association to be confirmed in Peru, ref.: García-Fuentes *et al*. 2020)

Forests flooded only by very high tides or storms, in soils with low saline concentration. Infratropical semiarid-arid. Biogeography: 2.1, 2.2.

###### CLASS XIII. BATIDO-SARCOCORNIETEA AMBIGUAE

Borhidi 1996, nom. mut. Galán,

Linares, Campos & Vicente 2009 [*Batido-Salicornietea ambiguae*]

+ Sarcocornietalia neei Galán, Linares, Campos & Vicente 2009

* Sarcocornio neei-Distichlion spicatae Galán, Linares, Campos & Vicente 2009

**111. *Cresso truxillensis-Distichlietum spicatae*** Galán, Linares, Campos & Vicente 2009 Vegetation on saline sandy soils. Thermotropical hyperarid. Biogeography: 3.3.

**112. *Cypero laevigati-Distichlietum spicatae*** Galán, Linares, Campos & Vicente 2009

Association on wet saline soils that do not become waterlogged. Thermotropical hyperarid. Biogeography: 3.3.

**113. *Sarcocornietum neei*** Müller & Gutte 1983, nom. mut. Galán, Linares, Campos & Vicente 2009 [*Salicornietum peruvianae*]

Salt marsh association with deep soils and intermittent flooding. Thermotropical hyperarid. Biogeography: 3.1, 3.3.

**114. *Sporobolo virginici-Distichlietum spicatae*** Galán, Linares, Campos & Vicente 2009

Coastal couch grassland bordering the littoral desert. Thermotropical hyperarid. Biogeography: 3.3.

##### G.2. VEGETATION OF INLAND SALINE SOILS

###### CLASS XIV. DISTICHLIO HUMILIS-ANTHOBRYETEA TRIANDRI

Navarro 1993

+ Anthobryetalia triandri Navarro 1993

* Sarcocornion pulvinatae Ruthsatz in Navarro 1993 [*Salicornio-Distichlion humilis* Ruthsatz 1977, nom. inval. Art. 3b (prov.)]

**115. *Distichlietum humilis*** Gutte & Müller 1985

Grassland in areas with saline efflorescence that dry out during the arid period. Thermo-supratropical arid-semiarid. Biogeography: 4.5.

**116. *Muhlenbergio fastigiatae-Distichlietum humilis*** Navarro 1993

Couch grasslands in high Andean areas with saline efflorescences, sometimes with sandy soils. Upper Supratropical and Orotropical arid-semiarid. Biogeography: 5.1.

**117. *Sarcocornietum andinae*** ass. nov. [*Salicornietum cuscoensis* Gutte & Müller 1985, nom. inval. Art. 3g, 3l]

Vegetation of endorheic basins with succulent cushion scrubs and grasses. Meso-supratropical semiarid-arid. Biogeography: 4.5. Characteristic plants: *Distichlis humilis, Sarcocornia andina, Spergularia marina.* Holotypus: *Sarcocornia andina* 4, *Distichlis humilis* +, *Spergularia marina* 2 (Elevation 3200 m, area 4 m^2^, Cuzco, Huarcapay, 19L 203845-8492646).

##### H. VEGETATION OF MARITIME DUNES

###### CLASS XV. IPOMOEO PEDIS-CAPRAE-TOURNEFORTIETEA GNAPHALODIS

Borhidi

1996, nom. mut. nov. [*Ipomeo pedis-caprae-Mallotonietea gnaphalodis*] [*Ipomoeo-Mallotonietea* Knapp 1964, nom. inval. Art. 7 & 8; *Canavalietea maritimae* Eskuche 1973, nom. inval. Art. 8]

+ Canavalio roseae-Ipomoeetalia pedis-caprae Borhidi 1996, nom. mut. nov. [*Canavalio maritimae-Ipomoeetalia pedís-caprae; Canavalio-Ipomoeetalia* Knapp 1964, nom. inval. Art. 7 & 8; *Suriano-Canavalietalia* Samek 1973, nom. inval. Art. 3b (prov.)]

* Ipomoeo pedis-caprae-Canavalion roseae Samek 1973, nom. mut. nov. [*Ipomoeo-Canavalion maritimae;* lectotypus: Borhidi 1996, p. 513]

**118. *Ipomoeo pedis-caprae-Canavalietum roseae*** Samek 1973, nom. mut. nov. [*Ipomoeo-Canavalietum maritimae;* lectotypus: Borhidi 1996, pp. 513-514] (association to be confirmed in Peru, ref. Eskuche 1973, Borhidi 1996)

Pioneer association of sandy beaches in northern Peru. Infratropical and Thermotropical, semiarid to hyperarid. Biogeography: 2.1, 2.2.

##### I. HERBACEOUS VEGETATION OF THE COASTAL HILLS OF THE PACIFIC DESERT

###### CLASS XVI. TILLANDSIETEA LATIFOLIO-LANDBECKII

cl. nov. [*Tillandsietea latifoliae* Müller 1985b, nom. inval. Art. 3b (prov.); *Tillandsietea landbeckii* Galán & Gómez 2003, nom. inval. Art. 17]

Plant communities of the sandy hills of the Pacific Desert, which behave like extreme xerophytes, with their root system atrophied, capturing atmospheric moisture through their leaves (aerophytes) (see Galán de Mera et al. 1999). They almost occur as monospecific groups, sometimes very dense and mostly located on mobile dunes. Characteristic plants: *Tillandsia landbeckii, T. latifolia, T. paleacea, T. purpurea, T. straminea, T. werdermannii.* Holotypus: *Tillandsietalia latifolio-landbeckii* ord. nov.

+ Tillandsietalia latifolio-landbeckii ord. nov. [*Tillandsietalia latifoliae* Müller 1985b, nom. inval. Art. 3b (prov.), *Tillandsietalia landbeckii* Galán & Gómez, nom inval. Art. 17]

This is the unique order known, with the same characteristic plants as the class. Holotypus: *Tillandsion purpureo latifoliae* all. nov.

* Tillandsion purpureo-latifoliae all. nov. [*Tillandsion latifoliae* Müller 1985b, nom. inval. Art. 3b (prov.); *Tillandsion latifoliae* Galán & Gómez 2003, nom. inval. Art. 17]

Alliance from central and northern Peru. Characteristic plants: *Tillandsia latifolia, T. paleacea, T. purpurea.* Holotypus: *Tillandsietum purpureo-latifoliae* Müller 1985b

**119. *Tillandsietum latifolio-paleaceae*** Müller 1985b

Aerophytes on somewhat stony clayey soil. Thermotropical hyperarid. Biogeography: 3.1.

**120. *Tillandsietum purpureo-latifoliae*** Müller 1985b [*Tillandsietum purpureo-latifoliae* Galán & Gómez 2003, nom. superfl. Art. 29c]

Aerophytes of stabilized sandy hills in central Peru. Thermotropical hyperarid-ultrahyperarid. Biogeography: 3.1, 3.2, 3.3.

* Tillandsion werdermannii Galán & Gómez 2003

**121. *Cistantho tovarii-Tillandsietum werdermannii*** Galán 1996

Association settled on the sandy plains of Moquegua and Tacna. Thermotropical hyperarid. Biogeography: 3.3.

###### CLASS XVII. PALAUO DISSECTAE-NOLANETEA GAYANAE

Galán 2005 [*Commelinetea*

*fasciculatae* Müller 1985b, nom. inval. Art. 3b (prov.); *Oxalidetea bulbigerae* Müller 1985b, nom. inval. Art. 3b (prov.); *Eragrostietea peruvianae* Müller 1985b, nom. inval. Art. 3b (prov.); *Heliotropetea arborescentis* Müller 1985b, nom. inval. Art. 3b (prov.); *Trixetea paradoxae* Müller 1985b, nom. inval. Art. 3b (prov.)]

+ Commelinetalia fasciculatae Galán & Rosa in Galán, Rosa & Cáceres 2002 [*Commelinetalia fasciculatae* Müller 1985b, nom. inval. Art. 3b (prov.); *Heliotropetalia arborescentis* Müller 1985b, nom. inval. Art. 3b (prov.); *Trixetalia paradoxae* Müller 1985b, nom. inval. Art. 3b (prov.)]

* Loasion urentis Galán & Rosa in Galán, Rosa & Cáceres 2002 [*Commelinion fasciculatae* Müller 1985b, nom. inval. Art. 3b (prov.); *Heliotropion arborescentis* Müller 1985b, nom. inval. Art. 3b (prov.); *Koelerion trachyanthae* Müller 1985b, nom. inval. Art. 3b (prov.)]

**122. *Hymenocallido amancais-Trixetum paradoxae*** Müller 1985b

Association of basal areas of rocky creeks and gravel-rich alluvial fans. Thermotropical hyperarid-arid. Biogeography: 3.1.

**123. *Loaso nitidae-Commelinetum fasciculatae*** Müller 1985b

Association that settles on coarse boulders at the base of the hills. Thermotropical hyperarid. Biogeography: 3.1.

**124. *Paspalo penicillati-Valerianetum chaerophylloidis*** Müller 1985b

Association of the upper parts of the hills with clayey and stony soils. Thermotropical hyperarid-arid. Biogeography: 3.1.

**125. *Philoglosso peruvianae-Urocarpidetum peruviani*** Galán & Rosa in Galán, Rosa & Cáceres 2002

Nitrophilous and grazed vegetation of the Pacific Desert hills. Thermotropical hyperarid. Biogeography: 2.2, 3.1, 3.3.

+ Tetragonio crystallinae-Plantaginetalia limensis Galán, Linares, Campos & Vicente 2011a [*Oxalidetalia bulbigerae* Müller 1985b, nom. inval. Art. 3b (prov.); *Eragrostietalia peruvianae* Müller 1985b, nom. inval. Art. 3b (prov.)]

* Nolanion humifusae Galán, Linares, Campos & Vicente 2011a [*Nolanion humifusae* Müller 1985b, nom. inval. Art. 3b (prov.); *Nolanion gayanae* Müller 1985b, nom. inval. Art. 3b (prov.)]

**126. *Calandrinio paniculatae-Nolanetum humifusae*** Müller 1985b

Pioneer vegetation on stony soils rich in sands. Thermotropical hyperarid-arid. Biogeography: 3.1.

**127. *Drymario weberbaueri-Nolanetum gayanae*** Müller 1985b

Pioneer vegetation in deep and humid soils of central Peru. Thermotropical hyperarid-arid. Biogeography: 3.1.

**128. *Palauo rhombifoliae-Nolanetum gayanae*** Müller 1985b

Association on consolidated sandy soils from central Peru. Thermotropical hyperarid. Biogeography: 3.1.

**129. *Tetragonio crystallinae-Nolanetum gayanae*** Müller 1985b

Association on poorly stabilized soils from the hills of central Peru. Thermotropical hyperarid. Biogeography: 3.1.

* Nolanion spathulatae Galán, Linares, Campos & Vicente 2011a

**130. *Hoffmannseggio mirandae-Palauetum weberbaueri*** Galán, Linares, Campos & Vicente 2009

Vegetation in consolidated hillside sands from southern Peru (Mejía area, Arequipa). Thermotropical hyperarid. Biogeography: 3.3.

**131. *Nolanetum scaposo-spathulatae*** Galán, Linares, Campos & Vicente 2011a

Vegetation on poorly stabilized soils from the hills of southern Peru (Camaná area, Arequipa). Thermotropical hyperarid. Biogeography: 3.3.

**132. *Nolano spathulatae-Palauetum dissectae*** Galán, Linares, Campos & Vicente 2009

Vegetation on poorly stabilized soils from the hills of southern Peru (Mejía area, Arequipa). Thermotropical hyperarid. Biogeography: 3.3.

**133. *Palauetum camanensis-weberbaueri*** Galán, Linares, Campos & Vicente 2011a

Vegetation on consolidated hillside sands from southern Peru (Camaná area, Arequipa). Thermotropical hyperarid. Biogeography: 3.3.

##### J. SHORT ANNUAL PIONEER VEGETATION

###### CLASS XVIII. CRASSULETEA CONNATAE

Galán 1999 [*Chondrosomo simplicis-Muhlenbergietea peruvianae* Rivas-Martínez & Navarro in Navarro & Maldonado 2002, syntax. syn.]

+ Crassuletalia connatae Galán 1999 [*Chondrosomo simplicis-Muhlenbergietalia peruvianae* Navarro in Navarro & Maldonado 2002, syntax. syn.; *Legousietalia biflorae* Müller 1985b, nom. inval. Art. 3b (prov.)]

* Peperomion hillii Galán 1999 [*Legousion biflorae* Müller 1985b, nom. inval. Art. 3b (prov.)]

**134. *Legousio biflorae-Centaurietum lomae*** Müller 1985b

Pioneer vegetation on clay soils in the coastal hills of the Peruvian desert. Thermotropical hyperarid-arid. Biogeography: 3.1.

**135. *Peperomio hillii-Crassuletum connatae*** Galán 1999

Pioneer vegetation on lithosols from the coastal hills of the Peruvian desert. Thermotropical hyperarid-arid. Biogeography: 3.1.

* Monnino pterocarpae-Cyperion andinae Galán, Linares, Campos, Trujillo & Vicente 2012

**136. *Hypseocharito bilobatae-Boutelouetum simplicis*** Galán, Linares, Campos, Trujillo & Vicente 2012

Pioneer therophytic vegetation on clay soils. Supratropical dry. Biogeography: 4.8.

**137. *Monnino ramosae-Boutelouetum simplicis*** Galán, Linares, Campos, Trujillo & Vicente 2012 Pioneer therophytic vegetation on acid sandy soils. Supratropical dry. Biogeography: 4.8.

**138. *Pectocaryo lateriflorae-Boutelouetum simplicis*** Galán, Linares, Campos, Trujillo & Vicente 2012

Pioneer therophytic vegetation on clay soils from western basal areas. Mesotropical semiarid. Biogeography: 5.1.

#### K. WETLAND VEGETATION

##### K.1. VEGETATION OF DEPRESSIONS, SPRINGS AND PEAT BOGS

###### CLASS XIX. MIMULO GLABRATI-PLANTAGINETEA AUSTRALIS

cl. nov. [*Plantaginetea australis* Gutte 1986, nom. inval. Art. 3b (prov.); *Plantaginetea australis* Gutte 1987a, nom. inval. Art. 8]

Class comprising plant communities of springs, small streams, and flat sloping peatlands of the Andean mountain range. Characteristic plants: *Calamagrostis rigescens, Eleocharis albibracteata, Epilobium denticulatum, Mimulus glabratus, Plantago australis, Polypogon interruptus.* Holotypus: *Mimulo glabrati-Plantaginetalia australis* ord. nov.

+ Mimulo glabrati-Plantaginetalia australis ord. nov. [*Marchantio-Epilobietalia* Cleef 1981, nom. inval. Art. 2d & 3g; *Marchantio plicatae-Epilobietalia denticulati* Cleef 1981 apud Cleef et al. 2008, nom. inval. Art. 6; *Plantaginetalia australis* Gutte 1987a, nom. inval. Art. 8]

Neotropical order. Characteristic plants: the same plants as in the class. Holotypus: *Mimulion glabrati* Galán, Linares, Campos, Trujillo & Vicente 2012

* Mimulion glabrati Galán, Linares, Campos, Trujillo & Vicente 2012 [*Plantaginion australis*

Gutte 1987a, nom. inval. Art. 8]

**139. *Calamagrostio spicigerae-Gentianelletum incurvae*** Gutte 1985

Association of humid depressions and springs located in central Peru. Orotropical humid. Biogeography: 4.3.

**140. *Mimulo glabrati-Polypogonetum interrupti*** Galán, Linares, Campos, Trujillo & Vicente 2012 [Comunidad de *Polypogon interruptus* y *Eleocharis geniculata* Galán, Cáceres & González 2003, nom. inval. Art. 3c; Comunidad de *Rorippa nasturtium-aquaticum* y *Veronica anagallis-aquatica* Galán, Cáceres & González 2003, nom. inval. Art. 3c]

Vegetation of streams and ditches with fast flowing waters. Thermo-supratropical, arid-subhumid. Biogeography: 5.1.

**141. *Nasturtio officinalis-Commelinetum communis*** Gutte 1978 [*Veronica anagallis-aquatica-Nasturtium officinale* Ges. Gutte 1986, nom. inval. Art. 3c, *Nasturtium officinale* Ges. Seibert & Menhofer 1991, nom. inval. Art. 3c]

Vegetation on streams and ditches with intermittent waters. Thermo-supratropical hyperarid-subhumid. Biogeography: 3.1.

**142. *Ranunculo praemorsi-Calamagrostietum rigescentis*** Gutte 1985

Association on humid plains suitable for grazing, from central Peru Supratropical humid. Biogeography: 4.3.

***Sisyrincho tinctorii-Plantaginetum australis*** Gutte 1986 [*Calamagrostio rigescentis-Plantaginetum australis* Gutte 1986, syntax. syn.]

Peat bogs settled near watercourses, sometimes with a scant slope, rich in clays and silts. Orotropical subhumid-humid. Biogeography: 4.3, 4.8.

###### CLASS XX. PLANTAGINI RIGIDAE-DISTICHIETEA MUSCOIDIS

Rivas-Martínez & Tovar

1982 [*Plantaginetea rigidae* Gutte 1980, nom. inval. Art. 2b & 8; *Wernerietea* Cleef 1981, nom. inval. Art. 3b (prov.)]

+ Calamagrostio jamesonii-Distichietalia muscoidis Rivas-Martínez & Tovar 1982

* Calamagrostio jamesonii-Distichion muscoidis Rivas-Martínez & Tovar 1982 [*Distichion muscoidis* Gutte 1980 p.p., nom. inval. Art. 3b (prov.)]

**143. *Calamagrostio jamesonii-Distichietum muscoidis*** Rivas-Martínez & Tovar 1982 [Asocies de *Distichia muscoides* y *Plantago rigida* Tovar 1973, p.p., nom. inval. Art. 3c]

Hard peatlands with slightly convex cushion vegetation. Orotropical subhumid-humid. Biogeography: 4.1, 4.3.

**144. *Isoeto andicolae-Distichietum muscoidis*** Gutte 1980, nom. mut. nov. [*Stylito andicolae-Distichietum muscoidis*]

Association at the margin of glacial lagoons, in the discontinuities of hard peat bogs. Oro-cryorotropical subhumid-humid. Biogeography: 4.1.

* Hypsello reniformis-Plantaginion rigidae Rivas-Martínez & Tovar 1982

**145. *Hypsello reniformis-Plantaginetum rigidae*** Rivas-Martínez & Tovar 1982 [*Carex microglochin-Plantago rigida* Ges. Gutte 1980, p.p., nom. inval. Art. 3c]

Association on flat and hard peat bogs of ponds, streams, and depressions, dominated by

*Plantago rigida* (‘champa estrella’). Orotropical subhumid-humid. Biogeography: 4.1, 4.3.

+ Calamagrostietalia nitidulae Galán, Cáceres & González 2003

* Calamagrostion chrysanthae Rivas-Martínez & Tovar 1982

**146. *Calamagrostietum nitidulo-chrysanthae*** Gutte 1980

Grassland causing the establishment of peat bogs. Orotropical dry-humid. Biogeography: 4.1, 5.1.

**147. *Calamagrostio ovatae-Wernerietum aretioidis*** Galán, Méndez, Linares, Campos & Vicente 2014

Cushion vegetation in glacier meltwater streams. Cryorotropical subhumid-humid. Biogeography: 4.1, 4.2.

**148. *Poo glaberrimae-Calamagrostietum eminentis*** ass. nov. [*Poo glaberrimae-Calamagrostietum eminentis* Rivas-Martínez & Tovar 1982, nom. inval. Art. 3b (prov.)]

Tall clumpy grassland at the edges of heavily drained lagoons and flowing streams. Orotropical subhumid-humid. Biogeography: 4.3. Characteristic plants: *Calamagrostis brevifolia, C. eminens, Poa glaberrima.* Holotypus: Rivas-Martínez & Tovar 1982: Tab. 4, plot 3.

+ Plantaginetalia tubulosae Gutte 1985 [lectotypus: Galán et al. 2003, p. 131]

* Hypsello reniformis-Plantaginion tubulosae Galán, Cáceres & González 2003

**149. *Eleocharito tucumanensis-Plantaginetum tubulosae*** Seibert 1993

Peatlands with cushion plants in depressions with oligotrophic waters. Oro-cryorotropical subhumid-humid. Biogeography: 4.7, 5.1.

* Oxychloion andinae Ruthsatz 1995

**150. *Wernerio pygmaeae-Puccinellietum oresigenae*** Galán, Cáceres & González 2003 Peat bogs with cushion plants in salty water. Orotropical dry. Biogeography: 4.8, 5.1.

##### K.2. HELOPHYTE VEGETATION

###### CLASS XXI. XYRIDO CAROLINIANAE-TYPHETEA DOMINGENSIS

1. O. Bolòs, Cervi &

Hatschbach 1991

+ Equiseto gigantei-Typhetalia domingensis O. Bolòs, Cervi & Hatschbach 1991

* Equiseto gigantei-Typhion domingensis O. Bolòs, Cervi & Hatschbach 1991

**151. *Bacopo monnieri-Typhetum domingensis*** Galán 1995

Tall helophytes of the neotropical Peruvian and Chilean coast. Infra-thermotropical hypearid. Biogeography: 2.1, 2.2, 3.1, 3.2, 3.3.

* Cortaderion jubatae Galán, Cáceres & González 2003

**152. *Cortaderietum jubatae*** Galán, Cáceres & González 2003

Tall helophytes of Andean watercourses. Meso-supratropical semiarid-humid. Biogeography: 5.1.

+ Oryzo grandiglumis-Hymenachnetalia amplexicaulis Galán & Rosa in Galán, Rosa & Cáceres 2002

* Hymenachnion amplexicaulis Galán 1995

**153. *Oxycaryo cubensis-Eleocharitetum acutangulae*** Galán & Linares 2008 (association to be confirmed in Peru, ref.: Galán 2014)

Vast floating masses of Cyperaceae, forming an herbaceous tapestry of *Oxycaryum cubense* and other species in the Amazonian Basin. Infratropical hyper-ultrahyperhumid. Biogeography: 7.1, 7.2, 8.1, 8.2.

**154. *Paspalo repentis-Pontederietum rotundifoliae*** Galán 1995

Association of floating grasses and water hyacinths with white flowers, characteristic of the oligotrophic waters of the Amazon Basin (‘barriales’, Ref. Amaz. Encarnación 1985, 1993). Infratropical ultra-hyperhumid. Biogeography: 7.2.

* Ipomoeion fistulosae Fuentes & Navarro 2000 [lectotypus: Galán 2014, p. 465]

**155. *Ipomoeo fistulosae-Sennetum aculeatae*** Galán & Linares 2008 (association to be confirmed in Peru, ref.: Galán 2014)

Amazonian hygro-nitrophile vegetation linked to fires. Infratropical hyper-ultrahyperhumid. Biogeography: 8.2.

+ Schoenoplectetalia olneyi-americani Galán, Linares, Campos & Vicente 2009

* Ludwigio octovalvis-Paspalion vaginati Galán, Linares, Campos & Vicente 2009

**156. *Lippio nodiflorae-Paspaletum vaginati*** Galán, Linares, Campos & Vicente 2009

*Paspalum* grassland from the Pacific coast, flooded by salt water. Thermotropical hyperarid. Biogeography: 3.3.

**157. *Samolo floribundi-Bacopetum monnierii*** Müller & Gutte 1983 [lectotypus hoc loco: Müller & Gutte 1983, tab. 5, plot 6]

Association on the edge of irrigation ditches. Thermotropical hyperarid. Biogeography: 3.1.

**158. *Schoenoplectetum olneyi-americani*** Galán, Linares, Campos & Vicente 2009 Pacific coast reed beds. Thermotropical hyperarid. Biogeography: 3.1, 3.2, 3.3.

* Schoenoplection tatorae all. nov.

Alliance that brings together the high Andean reed beds. Characteristic plants: *Cyperus sesslerioides, Juncus arcticus, J. cyperoides, Schoenoplectus tatora*. Holotypus: *Schoenoplectetum tatorae* Galán 1995

**159. *Schoenoplectetum tatorae*** Galán 1995, nom. mut. nov. [*Scirpetum tatorae*]

High Andean reed beds. Supra-orotropical dry-humid. Biogeography: 4.1, 4.2, 4.3, 4,7, 4.8.

+ Gynerio-Bambusetalia Borhidi 1996

* Gynerion sagittati Borhidi 1996

**160. *Tessario integrifoliae-Gynerietum sagittati*** Galán 1995

Shrub and rivercane association developing along Amazonian riversides and sandy beaches. Infratropical, hyper- and ultrahyperhumid. Biogeography: 7.1, 7.2, 8.1, 8.2.

* Montrichardion arborescentis Galán 1995

**161. *Montrichardietum arborescentis*** Galán 1995

Amazonian association of tall helophytes growing on abandoned meandering muds (‘pungales’, Ref. Amaz. Encarnación 1985, 1993). Infratropical, hyper- and ultrahyperhumid. Biogeography: 7.1, 7.2, 8.1, 8.2.

##### K.3. AMPHIBIOUS HIGH ANDEAN VEGETATION OF SMALL PLANTS

###### CLASS XXII. LIMOSELLETEA AUSTRALIS

cl. nov. [*Litorelleta australis* Oberdorfer 1960, nom. inval. Art. 3b (prov.); *Nanojuncetea autralis* Oberdorfer 1960, nom. inval. Art. 3b (prov.); Limoselletea Cleef 1981, nom. inval. Art. 3b (prov.); *Limoselletea* Cleef in Cleef, Rangel & Arellano 2008 nom. inval. Art. 3g & 8]

Amphibious vegetation consisting of therophytes and small perennial plants on the margins of lagoons and shallow-water ponds in the high Andean mountains and the southern Antarctic region. Characteristic plants: *Elatine triandra, Gratiola peruviana, Limosella australis, Ranunculus limoselloides, R. mandonianus.* Holotype: *Crassuletalia venezuelensis* Cleef 1981.

+ Crassuletalia venezuelensis Cleef 1981, nom. mut. Rangel & C. Ariza 2000 [*Tilleetalia*] [*Crassuletalia* Cleef in Cleef, Rangel & Arellano 2008, nom. superfl. Art. 29c]

* Ditricho submersi-Isoetion lechleri all. nov. [*Ditricho submersi-Isoetion* Cleef 1981 nom. inval. Art. 3g]

Neotropical Andean communities of small buttercups, bryophytes and isoetids submerged to a depth of about 1 m, on stony, pebbly, clayey, or peaty soils. Characteristic plants: *Blindia magellanica* (bryo.), *Ditrichum submersum* (bryo.), *Fissidens rigidulus* (bryo.), *Herbertus sendtneri* (bryo.), *Isoetes andicola, I. lechleri, Isotachis lacustris* (bryo.), *I. serrulata* (bryo.), *Riccardia cataractarum* (bryo.), *Sphagnum pylaesii.* Holotypus: *Isoetetum lechlerii* Cleef 1981.

**162. *Isoetetum lechlerii*** Cleef 1981, nom. mut. nov. [*Isoetetum sociae*] [*Isoetetum lechleri* Gutte 1980, nom. inval. Art. 5; *Isoetetum lechleri* Seibert 1993, syntax. syn.]

Association with isoetids forming a dark brownish-greenish carpet on the bed of glacial lakes with peaty soils. Orotropical, subhumid-humid. Biogeography: 4.1, 4.2, 4.3.

**163. *Ranunculetum limoselloidis*** Galán, Cáceres & González 2003

Aquatic vegetation of shallow oligotrophic water that leaves the interstices of the high Andean peat bogs from southern Peru. Orotropical subhumid-humid. Biogeography: 4.8, 5.1.

**164. *Ranunculetum mandoniani*** Galán 1995

Aquatic plant communities rooted at about 20 cm depth, in contact with high Andean peat bogs from Central Peru. Orotropical subhumid-humid. Biogeography: 4.2.

##### K.4. AMPHIBIOUS VEGETATION OF SMALL PLANTS FROM WARM AREAS

###### CLASS XXIII. XYRIDETEA SAVANENSIS

Galán 1995

+ Eleocharitetalia minimae Galán, González, Morales, Oltra & Vicente 2006

* Echinodorion boliviani Galán & Linares 2008

**165. *Bacopo myriophylloidis-Eleocharitetum minimae*** Galán & Linares 2008 (association to be confirmed in Peru, ref.: Galán 2014)

Flooded grassland rich in annual plants on silty soil. Infratropical, hyper- and ultrahyperhumid. Biogeography: 8.2.

**166. *Lindernio crustaceae-Xyridetum savanensis*** Galán 1995

Flooded grassland rich in annual plants on sandy soils. Infratropical, hyper- and ultrahyperhumid. Biogeography: 7.2.

##### K.5. FLOATING, TEMPORALLY ROOTED, AND NATANT LEAFY PLANT COMMUNITIES

###### CLASS XXIV. LEMNETEA MINORIS

1. R. Tx. ex O. Bolòs & Masclans 1955 [*Salvinio auriculatae-Eichhornietea crassipedis* Borhidi 1996, nom. amb. Art. 36]

+ Lemnetalia aequinoctialis Schwabe-Braun & R. Tx. 1981 nom. mut. Landolt 1986 [*Lemnetalia paucicostatae*]

* Azollo carolinianae-Salvinion auriculatae Borhidi 1996 [*Salvinio minimae-Lemnion aequinoctialis*

Landolt 1999, nom. inval. Art. 3b (prov.)]

**167. *Lemnetum aequinoctialis*** ass. nov. [*Lemno aequinoctialis-Wolffielletum lingulatae* Landolt 1999, nom. inval. Art. 3b (prov.)] (association to be confirmed in Peru, ref.: Landolt 1999, Galán 2014)

Duckweed communities of eutrophic waters in regions with warm winters. Infra-Mesotropical hyperarid to hyperhumid. Biogeography: 3.1, 7.2. Characteristic plants: *Azolla caroliniana, Lemna aequinoctialis, L. valdiviana, Ricciocarpus natans, Wolffiella lingulata, W. oblonga.* Holotypus: Landolt 1999: Tab. 2, plot 130.

* Azollo filiculoidis-Lemnion gibbae all. nov. [*Azollo filiculoidis-Lemnion gibbae* Landolt 1999, nom. inval. Art. 3b (prov.)]

Alliance of cold-water duckweed communities, mainly from high Andean areas and the Argentinean pampas with mild winters. Characteristic plants: *Azolla filiculoides, Lemna gibba, L. minuta, L. valdiviana.* Holotypus: *Lemnetum minuto-gibbae* Libermann Cruz, Pedrotti & Venanzoni 1988

**168. *Lemnetum minuto-gibbae*** Libermann Cruz, Pedrotti & Venanzoni 1988, nom. mut. Landolt 1999 [*Lemnetum minusculo-gibbae*]

Andean acropleustophyte vegetation of oligotrophic waters. Supra-orotropical semiarid to humid. Biogeography: 4.7, 5.1.

###### CLASS XXV. PISTIO STRATIOTIDIS-EICHHORNIETEA CRASSIPEDIS

1. O. Bolòs, Cervi & Hatschbach 1991 [*Eichhornietea crassipedis* Galán & Navarro 1992, syntax. syn.; *Salvinio auriculatae-Eichhornietea crassipedis* Borhidi 1996, p.p., nom. amb. Art. 36]

+ Pistio stratiotidis-Eichhornietalia crassipedis O. Bolòs, Cervi & Hatschbach 1991 [*Eichhornietalia crassipedis* Galán & Navarro 1992, syntax. syn.; *Salvinio auriculatae-Eichhornietalia crassipedis* Borhidi 1996, p.p., nom. amb. Art. 36]

* Pistio stratiotidis-Eichhornion crassipedis O. Bolòs, Cervi & Hatschbach 1991 [*Eichhornion crassipedis* Vu Van Cuong 1974, nom. inval. Art. 1; *Eichhornion crassipedis* Galán & Navarro 1992, syntax. syn.; *Eichhornion azureae* Borhidi 1996]

**169. *Eichhornietum azureae*** Borhidi in Borhidi, Muñiz & Del Risco 1983

Communities of *Pontederia azurea* in oligotrophic to mesotrophic deep freshwaters. Thermotropical humid-hyperhumid. Biogeography: 6.2.

**170. *Eichhornietum crassipedis*** Samek & Moncada 1971 [*Pistietum stratiotidis* Borhidi in Borhidi, Muñiz & Del Risco 1983, syntax. syn.]

Communities of *Pontederia crassipes* in eutrophic shallow waters. Thermotropical hyperarid-ultrahyperhumid. Biogeography: 3.1, 3.3, 7.2, 8.2.

###### CLASS XXVI. POTAMOGETONETEA

Klika in Klika & Novák 1941 [*Cabombo-Nymphaeetea* Borhidi 1996, nom. inval. Art. 3g]

+ Utricularietalia foliosae-junceae Borhidi 1996 [*Aldrovando-Utricularetalia* Borhidi in Borhidi, Muñiz & Del Risco 1983, nom. inval. Art. 3g]

* Utricularion foliosae-junceae Borhidi 1996 [*Aldrovando-Utricularion* Borhidi in Borhidi, Muñiz & Del Risco 1983, nom. inval. Art. 3g & 5]

**171. *Utricularietum foliosae*** Borhidi in Borhidi, Muñiz & Del Risco 1983

Free floating submerged water plant community. Infratropical hyperhumid to ultrahyperhumid. Biogeography: 7.2.

+ Nymphaeetalia amplae Borhidi 1996 [*Potamion striati* Müller & Gutte 1985, nom. inval. Art. 3b (prov.)]

* Myriophyllo quitensis-Potamion illinoensis Rangel & Aguirre 1983, nom. mut. nov. [*Myriophyllo elatinoidis-Potamion illinoensis*] [*Potamion illinoensis* Borhidi 1996, syntax. syn.]

**172. *Callitricho heteropodae-Alopecuretum hitchcockii*** Gutte 1987b

Association of floating leafy plants of stagnant or flowing clear waters from Central Peru. Orotropical subhumid-humid. Biogeography: 4.3.

**173. *Ceratophyllo demersi-Potamogetonetum striati*** Müller & Gutte 1985 [*Charo-Potametum striati* Müller & Gutte 1985, nom. inval. Art. 3g]

Submerged aquatic plant communities in the eutrophic flowing waters of irrigation ditches and canals. Themotropical hyperarid. Biogeography: 3.1.

**174. *Elodeetum potamogetonis*** Seibert 1993

Association of deep waters eutrophicated by nitrogen from livestock grazing in high Andean lagoons. Orotropical subhumid-humid. Biogeography: 4.3, 4.8, 5.1.

**175. *Hydrocotylo ranunculoidis-Myriophylletum aquatici*** Müller & Gutte 1985

Association of edges of irrigation ditches with eutrophic waters, with 20 to 50 cm depth. Thermotropical hyperarid. Biogeography: 3.1.

**176. *Myriophylletum quitensis*** Seibert 1993

Community of submerged plants in oligotrophic deep waters from high Andean lagoons. Orotropical subhumid-humid. Biogeography: 4.3, 4.8, 5.1.

**177. *Myriophyllo quitensis-Potamogetonetum illinoensis*** Rangel & Aguirre 1983, nom. mut. nov. Galán 1995 [*Myriophyllo elationoidis-Potametum illinoensis*] (association to be confirmed in Peru, ref.: Galán de Mera 1991)

A plant community of submerged and floating leafy plants of stagnant or slowly flowing dystrophic waters from northern Peru. Thermo-supratropical humid. Biogeography: 6.1, 6.2.

**178. *Stuckenietum punensis*** Galán, Cáceres & González 2003

Community of submerged plants in eutrophic flowing waters of high Andean streams and irrigation ditches. Supratropical subhumid. Biogeography: 4.3, 4.5, 4.7, 4.8.

* Nelumbo luteae-Nymphaeion amplae Samek & Moncada 1971

**179. *Nymphaeetum amplae*** Borhidi & Muñiz in Borhidi, Muñiz & Del Risco 1983

Floating leafy plant communities from deep, stagnant, dystrophic black waters of lakes and abandoned meanders in warm areas. Infratropical semiarid-ultrahyperhumid. Biogeography: 2.1, 8.2.

* Victorion amazonicae Galán 1995

**180. *Victorietum amazonicae*** Galán 1995

Vegetation of large nympheids from shallow, stagnant, eutrophic white waters of lakes and abandoned meanders in the Amazonia. Infratropical ultrahyperhumid. Biogeography: 7.2, 8.2.

##### L. ROCKY AND WALL VEGETATION

###### CLASS XXVII. DEUTEROCOHNIO LONGIPETALAE-PUYETEA FERRUGINEAE

Rivas & Navarro in Navarro & Maldonado 2002

+ Polypodio pycnocarpi-Puyetalia ferrugineae Galán & Rosa in Galán, Rosa & Cáceres 2002

* Peperomio galioidis-Puyion ferrugineae Galán & Rosa in Galán, Rosa & Cáceres 2002

**181. *Caricetum candicantis*** Galán, Baldeón, Beltrán, Benavente & Gómez 2004

Wild papaya bushes in mobile stony grounds and big boulder blocks. Thermo-mesotropical arid-dry. Biogeography: 4.1.

**182. *Matucano haynei-Tillandsietum humilis*** Galán, Baldeón, Beltrán, Benavente & Gómez 2004

Rupicolous vegetation very rich in xerophytes. Supratropical dry-subhumid. Biogeography: 4.1.

**183. *Polyachyro sphaerocephali-Puyetum densiflorae*** Galán, Linares, Campos & Vicente 2009

Association of large chasmophytes inhabiting wide fissures of vertical rock faces. Supratropical dry. Biogeography: 4.8.

###### CLASS XXVIII. WOODSIO MONTEVIDENSIS-CHEILANTHETEA PRUINATAE

cl. nov.

[*Notholaenetea niveae* Gutte 1986, nom. inval. Art. 3b (prov.), 5 & 8]

Neotropical vegetation of small rock crevices and walls with at least semi-arid rainfall interval, with abundant species of the genera *Cheilanthes* and *Asplenium*, and Polypodiaceae. Characteristic plants: *Argyrochosma nivea, Campyloneurum angustifolium, Cheilanthes pilosa, Ch. pruinata, Cystopteris fragilis, Leucheria daucifolia, Pleopeltis pycnocarpa, Saxifraga magellanica, Valeriana nivalis, Woodsia montevidensis.* Holotypus: *Salpichroetalia glandulosae* Galán, Cáceres & González 2003

+ Salpichroetalia glandulosae Galán, Cáceres & González 2003

* Belloo schultzii-Salpichroion glandulosae Galán, Cáceres & González 2003

**184. *Chersodomo diclinae-Valerianetum nivalis*** Galán, Cáceres & González 2003 [incl. *Belloo kunthianae-Brayopsietum calycinae* Chicalla 2017, nom. inval. Art. 5; Plantago sericeae-Gnaphalietum sp. Chicalla 2017, nom. inval. Art. 3b, 3g & 5]

Chasmophytic vegetation in andesitic rocks. Oro-cryorotropical dry-subhumid. Biogeography: 5.1.

* Woodsio montevidensis-Cheilanthion pruinatae Galán, Linares, Campos, Trujillo & Vicente 2012 [*Hypochaerido mucidae-Loricarion graveolentis* Montesinos-Tubée, Cleef & Sýkora 2015a, syntax. syn.]

**185. *Cheilanthetum arequipensis*** Galán, Linares, Campos, Trujillo & Vicente 2012 Chasmophytic vegetation on andesitic rocks. Mesotropical semiarid-dry. Biogeography: 5.1.

**186. *Oxalido petrophilae-Cheilanthetum pruinatae*** Galán, Linares, Campos, Trujillo & Vicente 2012

Chasmophytic vegetation on andesitic rocks. Supratropical dry-subhumid. Biogeography: 4.8.

**187. *Hypochaerido mucidae-Paronychietum muschleri*** Montesinos-Tubée, Cleef & Sýkora 2015a Chasmophytic vegetation on metamorphic rocks. Low Orotropical dry. Biogeography: 4.8.

**188. *Hypochaerido mucidae-Valeranietum nivalis*** Montesinos-Tubée, Cleef & Sýkora 2015a

Chasmophytic vegetation on andesitic rocks. Low Orotropical dry-subhumid. Biogeography: 4.8.

+ Saxifragetalia magellanicae Galán & Cáceres in Galán, Rosa & Cáceres 2002

* Saxifragion magellanicae Galán & Cáceres in Galán, Rosa & Cáceres 2002

**189. *Saxifrago magellanicae-Leucherietum daucifoliae*** Montesinos-Tubée, Cleef & Sýkora 2021

Chasmophytic association on poorly stable rocks formed by conglomerates with a high copper richness. Orotropical dry-subhumid. Biogeography: 4.8.

**190. *Valeriano thalictrioidis-Saxifragetum magellanicae*** Galán & Cáceres in Galán, Rosa & Cáceres 2002

Plant community form vertical rocks, generally of alkaline nature. Supra-orotropical subhumid-humid. Biogeography: 4.3.

###### CLASS XXIX. ADIANTETEA CAPILLI-VENERIS

Br.-Bl., Roussine & Nègre 1952 [*Oxalidetea megalorrhizae* Múller 1985 p.p., nom. inval. Art. 3b (prov.)]

+ Adiantetalia raddiani Galán & Rosa in Galán, Rosa & Cáceres 2002 [*Oxalidetalia megalorrhizae*

Müller 1985b p.p., nom. inval. Art. 3b (prov.)]

* Adiantion subvolubilis Galán & Rosa in Galán, Rosa & Cáceres 2002 [*Oxalidion megalorrhizae*

Müller 1985b, nom. inval. Art. 3b (prov.)]

**191. *Adiantetum subvolubilis*** Galán & Rosa in Galán, Rosa & Cáceres 2002 [*Pteris vittata-Adiantum capillus-veneris* Ges. Müller & Gutte 1985, nom. inval. Art. 3c]

Association of rock fissures and walls with oozing water in the coast of Ecuador and Peru. Infra-thermotropical hyperarid-arid. Biogeography: 3.1, 3.3.

**192. *Begonio geraniifoliae-Adiantetum subvolubilis*** Müller 1985b

Vegetation in slopes with loams and clays from the hills of the Peruvian desert. Thermotropical hyperarid-arid. Biogeography: 3.1.

**193. *Begonio octopetalae-Valerianetum pinnatifidae*** Müller 1985b

Vegetation of fissures in the upper parts of hills with oozing water. Thermotropical hyperarid-arid. 3.1.

#### M. VEGETATION OF ANTHROPIC ENVIRONMENTS

##### M.1. ANDEAN VEGETATION RUDERAL AND ADAPTED TO TRAMPLING

###### CLASS XXX. SONCHO-BIDENTETEA PILOSI

Hoff in Hoff, Brisse & Grandjouan 1983

+ Calandrinietalia ciliatae Galán, Linares, Campos, Trujillo & Vicente 2012 [*Capselletalia rubellae*

Gutte 1995, prov. Art.: 3b]

*Hordeion mutici Galán, Linares, Campos, Trujillo & Vicente 2012

**194. *Nassello pubiflorae-Stipetum mucronatae*** Galán, Linares, Campos, Trujillo & Vicente 2012

Vegetation linked to crops cultivated with Roman plough. Supra- and lower Orotropical subhumid. Biogeography: 4.8.

*Calandrinion ciliatae Gutte 1995

**195. *Calandrinietum ciliatae*** Seibert & Menhofer ex Seibert 1993

Vegetation linked to crops cultivated with Inca plough (‘taclla’). Supra-orotropical dry. Biogeography: 4.8.

+ Polygono hydropiperoidis-Rumicetalia cuneifolii Galán, Linares, Campos, Trujillo & Vicente 2012

* Rumicion cuneifolii Galán, Linares, Campos, Trujillo & Vicente 2012

**196. *Senecioni rudbeckiifolii-Rumicetum obtusifolii*** Gutte 1986

Association at the base of orchard walls and ditch separations, in wet soils. Meso-supratropical semiarid-subhumid. Biogeography: 4.3, 5.1.

###### CLASS XXXI. POLYGONO-POETEA ANNUAE

Rivas-Martínez 1975

+ Alternanthero pungentis-Cynodontetalia dactylonis ord. nov. [*Cynodonto-Pennisetalia clandestinae* Knapp 1965, nom. inval. Art. 8]

Vegetation adapted to trampling in the streets of villages and farmyards, from the tropical Andean mountain range. Characteristic plants: *Alternanthera pungens, Cynodon dactylon, Gamochaeta coarctata* (= *G. spicata*), *Guilleminea densa, Lepidium didymum, Paronychia andina, Solanum acaule.* Holotypus: *Lepidio chichicarae-Cynodontion dactylonis* all. nov.

* Lepidio chichicarae-Cynodontion dactylonis all. nov. [*Sporobolion minoris* Seibert & Menhofer ex Seibert 1993, nom. inval. Art. 3b (prov.)]

Currently, we recognize only one alliance with the same characteristic plants as the order. Holotypus: *Alternanthero pungentis-Cynodontetum dactylonis* Gutte 1978.

**197. *Alternanthero pungentis-Cynodontetum dactylonis*** Gutte 1978

Vegetation in paved soils. Meso-orotropical dry-subhumid. Biogeography: 4.8, 4.3, 6.3.

**198. *Cynodonto dactylonis-Pennisetetum clandestini*** Antezana, Barco & Navarro 2003 [*Pennisetum clandestinum* Ges. Seibert & Menhofer ex Seibert 1993, nom. inval. Art. 3c]

Vegetation in ridges separating orchards. Meso-supratropical arid. Biogeography: 4.8, 5.1.

**199. *Dichondro repentis-Cynodontetum dactylonis*** Gutte 1978

Vegetation adapted to trampling in warm areas. Thermotropical hyperarid. Biogeography: 3.1.

**200. *Gamochaeto spicatae-Lepidietum didymi*** Seibert & Menhofer ex Seibert 1993 corr. Gutte 1995 [*Gamochaeto spicatae-Lepidietum bipinnatifidi*] [*Poa annua-Lepidium bipinnatifidum* Ges. Seibert & Menhofer 1991, nom. inval. Art. 3c, *Solanum acaule* Ges. Seibert & Menhofer 1992, nom. inval. Art. 3c]

Vegetation adapted to animal trampling in corrals, livestock barns and streets of villages. Supratropical dry-subhumid. Biogeography: 4.8.

**201. *Lepidio chichicarae-Polygonetum avicularis*** Gutte 1986

Vegetation of roads and crowded streets, with plants of European origin. Supratropical dry-subhumid. Biogeography: 4.1.

##### M.2. RUDERAL AND ADAPTED TO TRAMPLING VEGETATION FROM THE AMAZONIA, AND IRRIGATED AREAS

###### CLASS XXXII. SIDO-STACHYTARPHETAETEA INDICAE

Hoff in Hoff, Brisse & Grandjouan 1983 [*Parthenio hysterophori-Dichanthietea annulati* Balátová-Tuláčková in Balátová-Tuláčková & García 1987, syntax. syn.]

+ Dichanthietalia annulati Balátová-Tuláčková in Balátová-Tuláčková & García 1987 [*Eleusinetalia indicae* Knapp 1957, nom. inval. Art. 3e & 8]

*Aristido adscensionis-Chloridion virgatae Galán & Rosa in Galán, Rosa & Cáceres 2002

**202. *Boerhavio caribeae-Sidetum paniculatae*** Gutte 1978

Road vegetation in warm areas. Thermotropical hyperarid. Biogeography: 3.1.

**203. *Cenchro echinati-Chloridetum virgatae*** Gutte 1978 [*Cynodon dactylon-Chloris radiata* Ges. Müller & Müller 1985, nom. inval. Art. 3c]

Vegetation linked to irrigated crops. Thermotropical hyperarid to subhumid. Biogeography: 2.2, 3.1, 6.3.

**204. *Eclipto albae-Paspaletum racemosi*** Gutte 1978

Vegetation on nitrogenized and humid soils from warm areas. Thermo-mesotropical hyperarid-arid. Biogeography: 3.1.

* Hyptido verticillatae-Paspalion conjugati Balátová-Tuláčková in Balátová-Tuláčková & García 1987 [*Wissadulo periplocifoliae-Cassion torae* Galán, González, Morales, Oltra & Vicente 2006, nom. inval. Art. 3b (prov.)]

**205. *Eclipto prostratae-Echinochloetum crus-pavonis*** ass. nov. [*Eclipta prostrata-Echinochloa crus-pavonis* Ges. Gutte & Müller 1989, nom. inval. Art. 3c]

Association at ditch edges and puddles with dirty water resulting from anthropozoogenic actions. Infratropical hyperhumid-ultrahyperhumid. Biogeography: 8.2. Characteristic plants: *Alternanthera sessilis, Eclipta prostrata, Echinochloa colonum, E. crus-pavonis.* Holotypus: Gutte & Müller 1989: Tab. 5, plot 5.

**206. *Euphorbio berteroanae-Cynodontetum dactylonis*** Balátová-Tuláčková in Balátová-Tuláčková & García 1987, nom. mut. nov. [*Chamaesyco berteroanae-Cynodontetum dactylonis*] [*Eleusine indica-Cynodon dactylon* Ges. Gutte & Müller 1989, nom. inval. Art. 3c]

Roadside vegetation of heavily trampled sites. Thermo-infratropical hyperarid to hyperhumid. Biogeography: 3.1, 8.2.

**207. *Sido glomeratae-Cassietum torae*** Castroviejo & López 1985 [*Amaranthus spinosus-Cassia tora* Ges. Gutte & Müller 1989, nom. inval. Art. 3c]

Annual nitrophilous Caribbean-Amazonian communities in areas with excessive livestock grazing, roadsides and near houses, where disturbed soils can observe. Infratropical hyperhumid-ultrahyperhumid. Biogeography: 8.2.

**208. *Sporobolo juncei-Paspaletum notati*** Gutte 1978, nom. mut. nov. [*Sporobolus poiretii-Paspaletum notati*]

Association of irrigated or trampled climatic meadows. Thermotropical hyperhumid. Biogeography: 3.1.

**209. *Sporoboletum indici*** Balátová-Tuláčková in Balátová-Tuláčková & García 1987 [*Paspalum notatum-Sporobolus indicus* Ges. Gutte & Müller 1989, nom. inval. Art. 3c]

Vegetation of park lawns and other moderately transited areas that are grazed several times a year. Infratropical hyperhumid-ultrahyperhumid. Biogeography: 8.2.

+ Sido acutae-Panicetalia maximi Hoff in Hoff, Brisse & Grandjouan 1983

* Stachytarpheto indicae-Panicion maximi Hoff in Hoff, Brisse & Grandjouan 1983

**210. *Stachytarpheto indicae-Panicetum maximi*** Hoff in Hoff, Brisse & Grandjouan 1983 [*Centrosemo virginianae-Panicetum maximi* Balátová-Tuláčková in Balátová-Tuláčková & García 1987, syntax. syn.; *Panicum maximum* Fluren Gutte & Müller 1989, nom. inval. Art. 3c]

Pantropical association in dense corridors of tall perennial grasses very frequent on roadsides and old abandoned terrains. Infratropical hyperhumid-ultrahyperhumid. Biogeography: 8.2.

##### M.3. VEGETATION OF DISTURBED SOILS AND DUMPS

###### CLASS XXXIII. NICOTIANO GLUTINOSAE-AMBROSIETEA ARBORESCENTIS

Galán & Cáceres in Galán, Rosa & Cáceres 2002

+ Nicotianetalia paniculato-glutinosae Galán & Cáceres in Galán, Rosa & Cáceres 2002 [*Urocarpidetalia peruvianae* Müller 1985b, nom. inval. Art. 3b (prov.)]

* Jungion axillaris Galán, Baldeón, Beltrán, Benavente & Gómez 2004

**211. *Jungietum axillaris*** Galán, Baldeón, Beltrán, Benavente & Gómez 2004

Nitrophilous association with woody plants of stockyard walls, waste dumps and road slopes. Thermo-mesotropical hyperarid-dry. Biogeography: 4.1.

* Sicyo baderoae-Urticion magellanicae Galán & Cáceres in Galán, Rosa & Cáceres 2002 [*Urocarpidion peruvianae* Müller 1985b, nom. inval. Art. 3b (prov.)]

**212. *Amarantho gracilis-Chenopodietum muralis*** Gutte 1978

Hypernitrophilic vegetation with *Chenopodium*. Thermotropical hyperarid-arid. Biogeography: 3.1.

**213. *Flaverio bidentis-Chenopodietum muralis*** Gutte 1978

Hypernitrophilic vegetation with *Chenopodium*. Thermotropical dry. Biogeography: 6.3.

**214. *Nicotiano paticulatae-Urocarpidetum peruviani*** Müller 1985b

Vegetation of disturbed soils of the Peruvian desert. Thermotropical, hyperarid-arid. Biogeography: 3.1.

**215. *Urtico flabellatae-Caiophoretum sepiariae* Gutte 1986**

Nitrophilous vegetation of stockyard walls and rock hollows. Supratropical dry-subhumid. Biogeography: 4.1.

**216. *Urtico flavellatae-Urocarpidetum peruviani*** Galán, Linares, Campos, Trujillo & Vicente 2012

Vegetation of waste dumps, rubbish dumps, and disturbed soils. Meso-supratropical dry-subhumid. Biogeography: 4.8.

##### N. SAVANNAS

###### CLASS XXXIV. RHYNCHOSPORO CILIATAE-ERAGROSTIETEA MAYPURENSIS

Galán 2014

[*Leptocoryphio-Trachypogonetea* Van Donselaar 1965, p.p., nom. inval. Art. 5]

+ Trachypogonetalia spicati Van Donselaar 1965, nom. mut. Galán 2014 [*Trachypogonetalia plumosi*] [lectotypus: Galán 2014, p. 453]

* Rhynchosporo barbatae-Trachypogonion spicati Van Donselaar 1965, nom. mut. Galán 2014 [*Rhynchosporo barbatae-Trachypogonion plumosi*] [lectotypus: Galán 2014, p. 454]

**217. *Schizachyrio brevifolii-Paspaletum foliiformis*** Susach Campalans 1989, nom. mut. nov. [Schizachyrio brevifolii-Thrasietum petrosae] [lectotypus: Galán 2014, p. 455] (Association to be confirmed in Peru, ref. Tovar 1993)

Lateritic hard substrate savannas with *Curatella americana*. Thermotropical subhumid-humid. Biogeography: 6.3.

**218. *Schizachyrietum teneri*** Susach Campalans 1989 [lectotypus: Galán 2014, p. 455] (Association to be confirmed in Peru, ref. Tovar 1993)

Savanna association on gravel and rocky soils. Thermotropical subhumid. Biogeography: 6.2, 6.3.

### Final remark

This syntaxonomic scheme is an update of the previous one from 2002 (Galán de Mera et al. 2002), and of the 2005 and 2006 approaches (Galán de Mera 2005, Galán de Mera & Vicente Orellana 2006). In the future, we will continue adding units from the different Peruvian territories while more plots are taken.

## Acknowledgement

We would like to thank Brian Crilly for his editorial assistance.

